# Mechanism of defibrillation of cardiac tissue by periodic low-energy pacing

**DOI:** 10.1101/2023.03.16.533010

**Authors:** Pavel Buran, Thomas Niedermayer, Markus Bär

## Abstract

Rotating excitation waves and electrical turbulence in excitable cardiac tissue are associated with arrhythmias such as life-threatening ventricular fibrillation. Experimental studies (S. Luther et al., **Nature** 475, 235-239 (2011)). have shown that a time-periodic sequence of low-energy electrical far-field pulses is able to terminate fibrillation more gently than a single high-energy pulse. During this so called low-energy antifibrillation pacing (LEAP), only tissue near sufficiently large conduction heterogeneities, such as large coronary arteries, is activated. Based on extensive simulations and simple theoretical reasoning, we present a comprehensive unified mechanism for successful LEAP in two spatial dimensional systems, which is able to explain both the termination of stable spirals and of spatiotemporal chaos. We carried out extensive simulations (more than 500000 runs for each considered model) varying pacing periods, pacing field strength and initial conditions using a model of cardiac tissue perforated by blood vessels, which was found earlier to reproduce the behavior seen in the LEAP experiments for different dynamical regimes and different cellular models (P. Buran et al., **Chaos** 27, 113110 (2017) and **New J. Phys**. 24 083024 (2022)). We studied altogether three different cellular models to capture qualitatively different kinds of fibrillatory states like stable spirals and spatiotemporal chaos. To achieve a mechanistic understanding of the simulation results, we have investigated a variety of macroscopic observables characterizing an excitable medium with respect to their correlation with the success of an individual low-energy pulse during LEAP. We found in all considered cases that the refractory boundary length *L*_*RB*_, the total length of the borders between refractory and excitable parts of the tissue, displays the strongest correlation with the success of the pacing and thus predicts best the success of an individual LEAP pulse. Furthermore, we found the success probability *P*_*L*_ decays exponentially with this length according to *P*_*L*_ = *exp*(− *k*(*E*)*L*_*RB*_), where *E* is the strength of the electrical field in pacing and *k*(*E*) is a monotonically decreasing function of *E*. A closer look at the spatiotemporal dynamics in the simulations reveals that actually each pulse during LEAP annihilates practically all defects and excitation fronts, however, also induces new pairs of defects and associated excitation fronts at the refractory boundaries. The success probability of each individual pulse can thus be simply interpreted as the probability that no new rotor pair gets created by the shock, while all existing defects get annihilated. This assumption allows to derive the observed exponential dependence of the success probability on the refractory boundary length, where the prefactor *k*(*E*) in the exponent is equal (for stable spirals) or proportional (for spatiotemporal chaos) to the probability *λ*(*E*) that a new rotor pairs is created by the shock along a segment of unit length along the refractory boundary. Our findings are in conformity with the upper limit of vulnerability (ULV) hypothesis, which states that the single pulse defibrillation threshold is simply given by the lowest field strength, where no new rotor pairs arise as a result of the shock. LEAP operates at field strengths (and energies) below this ULV limit. Successful LEAP protocols are characterized by a coordinated interplay between the pulses, that gradually decreases the refractory boundary length and therefore simultaneously increases the success probability until complete defibrillation is achieved.

## I. INTRODUCTION

A loss of rhythm and synchronization of the cardiac electrical activity, orchestrating heart contraction and the pumping of blood, is associated with a number of arrhythmias including atrial fibrillation (AF) and ventricular fibrillation (VF). VF is a particularly dangerous malfunction of the heart, preventing an effective mechanical contraction of the ventricles and causing sudden cardiac death within a few minutes if left untreated. The electrical activity during VF is dominated by multiple stable spirals, one or a few large spirals surrounded by irregular activity or entirely by multiple wavelets in a spatiotemporally chaotic state.^1–4^. The nonlinear dynamics behind the emergence of complex dynamical states has been studied extensively, for review see e. g.^5–7^. Spontaneous termination of complex activity^8–10^ or the suppression of such dynamics by appropriate external forcing protocols in experiments^6,11–13^ and simulations^14–18^ are more recently tackled topics of interest.

Defibrillation by a strong electrical pulse is in practice the only known effective way of terminating VF. However, strong shocks are associated with adverse side effects including pain and trauma of the patient^19^ as well as damage of the myocardium.^20^ These adverse effects could be avoided or at least diminished if VF would be terminated reliably by defibrillation shocks of significantly lower energy. Recent experimental studies^12,13^ of AF *in vitro and in vivo* and VF *in vitro* have shown that a sequence of 5 low-energy far-field pulses with stimulation rates close to the arrhythmia cycle length can require 80% − 90% less energy per pulse than a single defibrillation shock. This method has been called low-energy antifibrillation pacing (LEAP). A similar energy reduction was also found with much faster pacing rates than the cycle length for AF,^21,22^ while the energy may be reduced further by an order of magnitude for ventricular tachycardia (VT).^23,24^

LEAP is based on the fact that an electric field depolarizes and hyperpolarizes the tissue close to conductivity heterogeneities^25^ which become virtual electrodes.^26^ The strength of this effect depends both on the strength of the electrical field and on the size and shape of the heterogeneities.^27–29^ Only tissue at mayor conduction heterogeneities may be activated by weak pulses. Therefore, global tissue activation and wave termination often originate only from few localized activation sites (hot spots). Luther *et al*.^13^ suggested that these hot spots are given by large coronary arteries and show that the size distribution of the heterogeneities follows a power law. Caldwell *et al*.^30^, however, emphasize that they did not find such a co-localization of hot spots and major coronary vessels.

The mechanism how LEAP terminate arrhythmias is not well understood yet since experimental methods to visualize the whole three dimensional electrical activity inside the heart are missing. A new promising method which might give deeper insights in the future is the determination of the three dimensional electrical activity over a ultrasound measurement of the mechanical activity of the heart.^31^ A common interpretation of LEAP is synchronization, suggested by the observations in experiments^12,13,32^ and simulations^32,33^ that usually each pulse gradually entrains more tissue during successful LEAP until the entire tissue is synchronized and no excitation waves are left. Current ths.comeoretical approaches to understand LEAP often focus on the process of unpinning and removal of a small number of stable spirals and suggest LEAP protocols using overdrive or underdrive pacing or combinations of it.^11,27,34,35^ In these papers, it was demonstrated that spirals that were pinned at larger heterogeneities are best terminated if the LEAP pacing frequency is set to 80% − 90% of the rotation frequency of the spiral around the pinning site. This choice allows for an efficient scanning of the phase of the spiral in order to find the vulnerable window for spiral termination.^15,36^

In a previous statistical study^14^ we have analyzed in detail the defibrillation success of LEAP in terms of the field strength, pacing period and number of pulses within a rather simple two dimensional model. Non-conducting heterogeneities were represented by circles whose size distribution is given by the distribution of radii of coronary arteries measured in Ref^13^. Two alternative cellular models^37,38^ were studied in detail which show different kinds of electrical turbulence and scaled such, that they exhibit similar length scales and the correct magnitude of the electric field strength necessary for single pulse defibrillation. In both cases, a resonant pacing with the dominant period of electrical turbulence resulted in maximal energy reduction and if the necessary field strength for LEAP was overestimated, underdrive pacing was superior to resonant pacing. However, the process of unpinning or a drift of spirals to the boundary was not observable. Also not in a following statistical study^17^, focusing on the termination of stable spirals and using a modified version of the Luo-Rudy model^39^. We could not even observe, studying the differences in the termination of either single free or pinned spirals as well as multiple free spirals, that the number of spirals or the fact whether a spiral is pinned or not has any essential impact on the success of LEAP. Instead we observed a progressively increase of the right before each pulse excitable tissue fraction during successful LEAP corresponding with the idea of synchronization. However, whether this increase is causal for the defibrillation success or only a side effect of LEAP, is not clarified yet.

The aim of the present work is to reveal and understand the basic mechanism behind LEAP. Therefore, we examine a wide range of macro-observables like the number of defects or the fraction of excitable tissue and their ability to possibly predict the success of LEAP. It is a continuation of our two above already outlined studies^14,17^ and uses for the simulations the same simple two dimensional model of cardiac tissue perforated by blood vessels for the three different in this two studies investigated cellular models, which exhibit different kinds of fibrillatory states. It turns out in all three cases that the refractory boundary length, the total length of the borders between refractory and excitable parts of the tissue, predicts best the success of each individual LEAP pulse and that the success probability decays exponentially with this length. Moreover we will show that actually each pulse during LEAP annihilates all excitation fronts, however, alo induces new excitation fronts at these borders between refractory and excitable parts of the tissue and that the success probability of each individual pulse can thus be simply interpreted as the probability that no new excitation front gets induced that arouses new fibrillation. In behalf of this, we will then derive the observed exponential dependence between the success probability and the refractory boundary length. Finally we will show how successful LEAP protocols achieve a gradual decrease of this length that is needed for successful defibrillation.

In Sec. II, we introduce the parameters and the cellular model used in a mono-domain framework, our choice of initial conditions and our selection of macro-observables. Sec. III contains the main results regarding the analysis of the impact of the macro-observables on the termination success, the derivation of the exponential relation between the termination probability and the refractory boundary length and the operation principle of successful LEAP protocols. Finally, Sec. IV provides a discussion of the results.

## II. METHODS

### A. Numerical methods and model parameters

This work is a continuation of our previous numerical studies about LEAP of electrical turbulence^14^ and stable spirals^17^. We used the same simple model of homogenous, isotropic and two-dimensional tissue which is perforated by blood vessels and which bases on the mono-domain model^40^ in conjugation with a cellular model. Furthermore we investigated the mechanism behind LEAP for all the three in this two studies tested cellular models, which exhibit different kinds of fibrillatory states, see Fig.1:

**FIG. 1.**
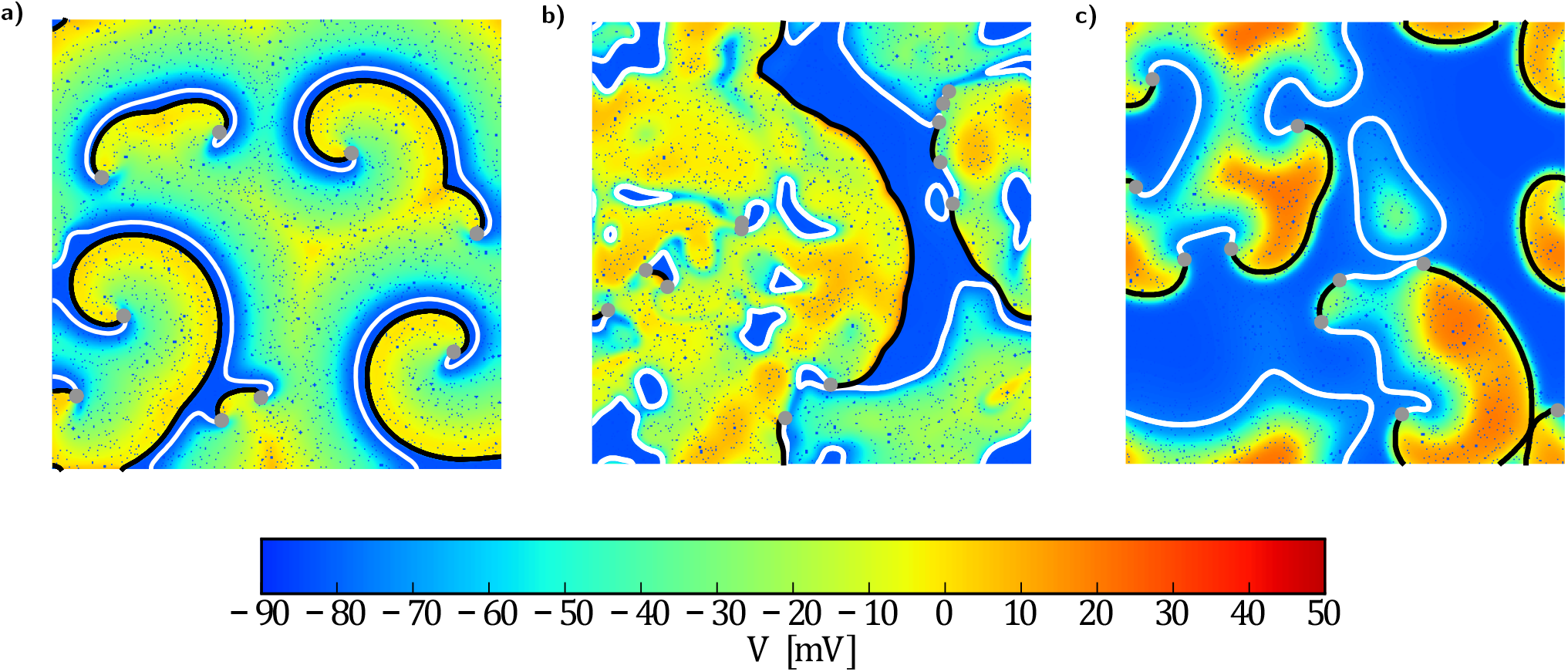
Snapshot of transmembrane potential *V* for the (a) (LRmod) model exhibiting stable spirals, (b) (LR) model exhibiting spatiotemporal chaos and (c) (FK) model, exhibiting unstable spirals. Conductivity heterogeneities are represented by small blue circles, excitation fronts are plotted black, refractory boundaries are plotted white and the topological defects are marked as grey dots. In all three models an average of 10 phase defects are found for the unperturbed states. Movies of the simulations are provided in the supplementary material.

a. A modified version of the Luo-Rudy model^37^ (LRmod) with parameter values from Ref^39^, exhibiting stable spirals.
b. The Luo-Rudy model^37^ (LR) with standard parameter values, exhibiting spatiotemporal chaos.
c. The Fenton-Karma model^38^ (FK) with *V*_0_ = − 84 mV, *V*_fi_ = 23.125 mV and default parameters from Ref^41^, exhibiting unstable spirals.

In the following, we will give only a brief overview about the used numerical methods, model parameters, distribution of conductivity heterogeneities and initial conditions, since these were adopted unchanged from our two previous studies. Further details about the (LR-mod) model can be found in the work for stable spirals^40^ and further details about the (LR) and (FK) model can be found in previous work for the case of electrical turbulence^14^.

All simulations were performed on a quadratic, 2-dimensional equidistant finite-difference grid with 1000 × 1000 nodess, resulting in a grid spacing of Δ*x* = 0.001 *L*, where *L* is the system size. No-flux boundary conditions were used for the (LRmod) model and periodic boundary conditions were used for the (LR) and (FK) model. The diffusion constants *D*_LRmod_ = 5.625 × 10^−3^ *L*^2^*/*s, *D*_LR_ = 2.5 × 10^−3^ *L*^2^*/*s and *D*_FK_ = 4.0 × 10^−2^ *L*^2^*/*s, were chosen such, that these models exhibit in average 10 phase defects, see Fig.1.

Conductivity heterogeneities were included in this grid as non-conducting patches by modified Neumann-boundary-conditions.^27,29^ and the mayor conducting heterogeneities were assumed to be given by large coronary arteries as suggested in Luther *et al*.^13^. They were represented by circles whose size distribution is given by the distribution of radii of this arteries and follows a power law^13^. The cutoffs of this size distribution *R*_min_ = 3 × 10^−4^ *L* and *R*_max_ = 4 × 10^−3^ *L* and the heterogeneity density *ρ*_0_ = 1.6 × 10^−4^ */L*^2^ were chosen such, that this choice resulted for all three models in single pulse defibrillation thresholds which were consistent with the one observed in experiments^42^. For all simulations the same realization of radii and positions were used, since different realizations of radii and positions result in almost identical defibrillation successes.

The time integration, were performed with a time step of 0.1 ms for the (LRmod) and (LR) model and 0.025 ms for the (FK) model, which are high enough in order to guarantee numerical stability.

For the (LRmod) model we have considered only the 50 initial conditions for the multiple spiral states, since these are the closest in representing a fibrillatory state and moreover, neither the number of spirals nor the fact whether a spiral is pinned or not had a significant impact on the success of LEAP for this model. For the (LR) and (FK) model we have simply taken the first 50 out of the 100 initial conditions.

### B. Defibrillation protocols

State-of-the-art defibrillators execute a biphasic, asymmetric protocol.^43^ Our LEAP protocol contains *n* identical biphasic pulses, where the interval between the onsets of two subsequent pulses is constant and given by a pacing period *T*. The waveform of the identical biphasic pulses consists of two subsequent rectangular parts with identical amplitudes *E*, opposite field direction and a duration of 7 ms (forward) and 3 ms (backward). If fixed to 10 ms, this configuration for the durations of the two phases led to best defibrillation results for all three in this work used cellular models.^14,17^

### C. Macro-observables in excitable media

The following 11 macro-observables were determined and analyzed in this work:

#### Fraction of excitable F_Exc_

Total fraction of excitable tissue (blue colored tissue in Fig.1).

#### Refractory boundary length L_RB_

Total length of all borders between refractory and excitable parts of the tissue (white lines in Fig.1).

#### Front length L_Fr_

Total length of all excitation fronts (black lines in Fig.1).

#### *∂*_t_F_Exc_

Time derivative of fraction of excitable

#### *∂*_*t*_L_RB_

Time derivative of refractory boundary length

#### *∂*_*t*_L_Fr_

Time derivative of the front length

#### Number of excitable clusters N_cExc_

Total number of all clusters consisting of excitable tissue (blue areas in Fig.1).

#### Number of non-excitable clusters N_cNExc_

Total number of all clusters consisting of non-excitable tissue (non-blue areas in Fig.1).

#### Number of refractory boundaries N_RB_

Total number of all borders between refractory and excitable parts of the tissue (white lines in Fig.1).

#### Number of fronts N_Fr_

Total number of all excitation fronts (black lines in Fig.1).

#### Number of defects N_Def_

Total number of all phase defects (grey dots in Fig.1).

Details about the determination of these macro-observables and especially about the segmentation of the refractory boundaries, excitation fronts and phase defects can be found in Appendix B 1.

### D. Simulation protocol

We have performed for all three cellular models LEAP simulations with 50 different initial conditions, 20 different pacing periods *T*, up to *n* = 10 pulses and different electric field strengths *E* following the recipes outlined in^14,17^. The field strengths *E* were chosen such, that they were spaced with 0.25 mV*/*cm equidistant between 0.0 mV*/*cm and the corresponding single pulse defibrillation threshold of the cellular model, given by 6.0 mV*/*cm for the (LRmod) model, 5.0 mV*/*cm for the (LR) model and 3.5 mV*/*cm for the (FK) model. The 20 chosen pacing periods *T* were spaced equidistant as well. They had values between 80 ms and 175 ms for the (LRmod) model, between 260 ms and 450 ms for the (LR) model and between 65 ms and 160 ms for the (FK) model. This choice of pacing periods ensures that a wide range of qualitative different dynamical states were considered for every field strength, since it covers the entire region of successful LEAP protocols as well as parts where LEAP is not successful, see Fig.2. In total we simulated for every field strength *E* the application of a LEAP pulse on 50 · 20 · 10 = 10000 different dynamical states. For all this simulations the defibrillation success *S* (“1” if the fibrillatory state is converted into a quiescent state by the pulse and “0” if not) and the eleven in Sec II C introduced macro-observables were determined.

**FIG. 2.**
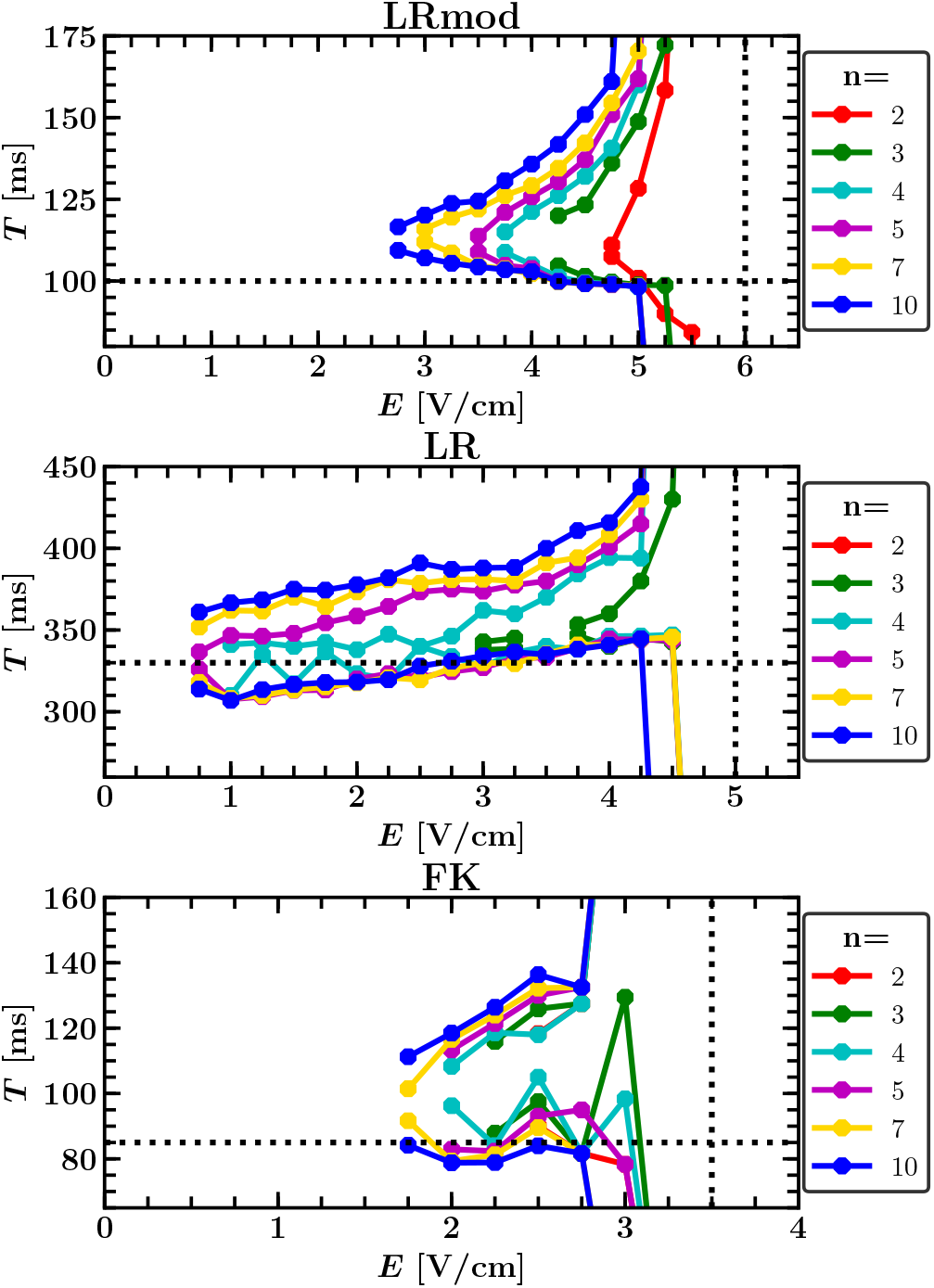
Borderlines of the regions of successful LEAP protocols for different pulse numbers *n*. This regions are defined as the range of pacing periods *T* and electric field strength *E*, where LEAP protocols with *n* pulses have a termination probability of at least 80%. These termination probabilities were computed for every parameter configuration as the fraction of successful termination events out of 50 simulation runs with different initial conditions. The vertical dotted lines mark the single pulse defibrillation thresholds, determined as the lowest electric field strength *E* with a single pulse termination probability of at least 80%. The dominant periods of the three models are marked by horizontal dotted lines.

## III. RESULTS

### A. Macro-observables and success rate of fibrillation by periodic pacing

The pacing period *T* was found to be crucial for LEAP success of both electrical turbulence^14^ and stable spirals^17^. Hence, LEAP cannot been explained as a combined effect of individual pulses. A certain degree of cooperativity of the pulses is therefore essential and there has to be a qualitative difference between the dynamical state of the system right before the first and right before the last pulse during successful LEAP, see Fig.3. The characterization of dynamical states which are susceptible to a single low-energy pulse is hence pivotal for understanding the cooperative nature of LEAP. To do so, the first objective was to correlate macro-observables that characterize a dynamical state by a single value and the the probability of a pulse to successfully terminate fibrillation in the medium *P*. For excitable media like cardiac tissue, we can qualitatively distinguish a rest state, an excited state and a refractory state, see e. g.^44^ or^6^. In addition, the dynamic states of an excitable medium is often characterized by the number of topological defects, that correspond to rotating stable spirals or rotating excitation in a chaotic state depending on the qualitative nature of the dynamics. Hence, we can define macro-variables such as the amount of excitable, excited or refractory tissues as well as the lengths of interfaces between connected areas of such states such as the excitation front (=interface between excitable and excited areas) or the refractory boundary (interface between refractory and excitable areas). Some features of dynamics in excitable media were also captured by discrete three-state cellular automata defining rules for the excitable, excited and refractory states and the transition between them^45^.

**FIG. 3.**
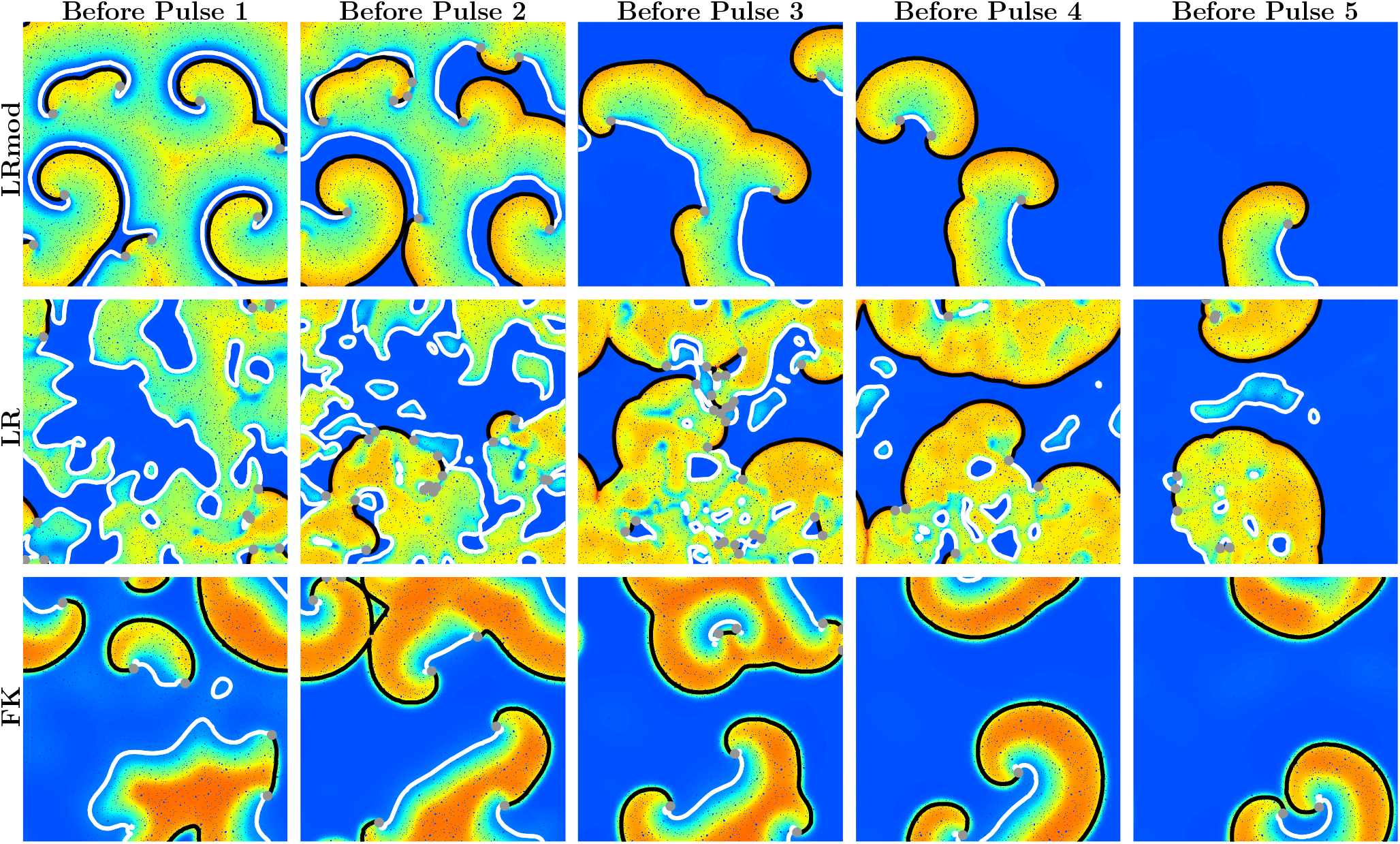
Snapshots of transmembrane potential right before the first, second, third, fourth and fifth pulse during LEAP with five pulses, where the first, second, third and fourth pulse do not terminate the fibrillation while the fifth does. What is the qualitative difference between the state right before the first pulse and right before the previous pulses and what is the characteristics which result in successful defibrillation? The applied LEAP protocols had an electric field strength of *E* = 3.5 mV*/*cm and pacing period of *T* = 115 ms for the (LRmod) model (left), a field strength of *E* = 1.0 mV*/*cm and pacing period of *T* = 340 ms for the (LR) model (center) and a field strength of *E* = 2.0 mV*/*cm and pacing period of *T* = 105 ms for the (FK) model (right). Movies of the simulations are provided in the supplementary material.

Ideally, the relationship between such a macro-observable *O* and the termination probability *P* should be describable by a simple monotonic function. A good measure for the monotonic relationships between two variables is Spearman’s rank correlation coefficient which has the advantage that it is robust against outliers and assesses even nonlinear relationships compared to Pearsons correlation coefficient which is normally used and assesses only linear relationships.^46^ Spearman’s rank correlation coefficient is appropriate for continuous and ordinal variables like all in Sec II C introduced macro-observables. However, the termination probability *P*, does not meet this requirements, even though it is continuous, it can be measured only dichotomously as the defibrillation success *S* which is defined as being “1” if a state got terminated and “0” if not. In this case, the underlying Spearman’s rank correlation coefficient can be still estimated by the rank biserial correlation^47^:

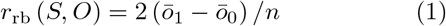

where *n* is the total sample size, *ō*_1_ and *ō*_0_ are the mean ranks of the continuous or ordinal macro-observable *O* for data pairs with a value of *s* = 1 and *s* = 0 respectively for the defibrillation success *S*.

We tested all eleven in Sec II C introduced macro-observables, for all three models and in each case for four different field strength *E*. The absolute values of the calculated Rank biserial correlation coefficients *r*_rb_ with the defibrillation success *S* are plotted in Fig. 4 for the refractory boundary length *L*_RB_, the front length *L*_Fr_, the fraction of excitable *F*_Exc_ and the number of defects *N*_Def_. All values of the calculated Rank biserial correlation coefficients *r*_rb_ for the remaining macro-observables are listed in Table I in Appendix B 2. The strongest correlations with the defibrillation success *S* were found for the refractory boundary length *L*_RB_. The refractory boundary length *L*_RB_ is furthermore the only macro-observable at all that is most strongly correlated with *S* for all three models and therefore the most promising macro-observable to identify a mechanism of LEAP that holds for all three models.

**FIG. 4.**
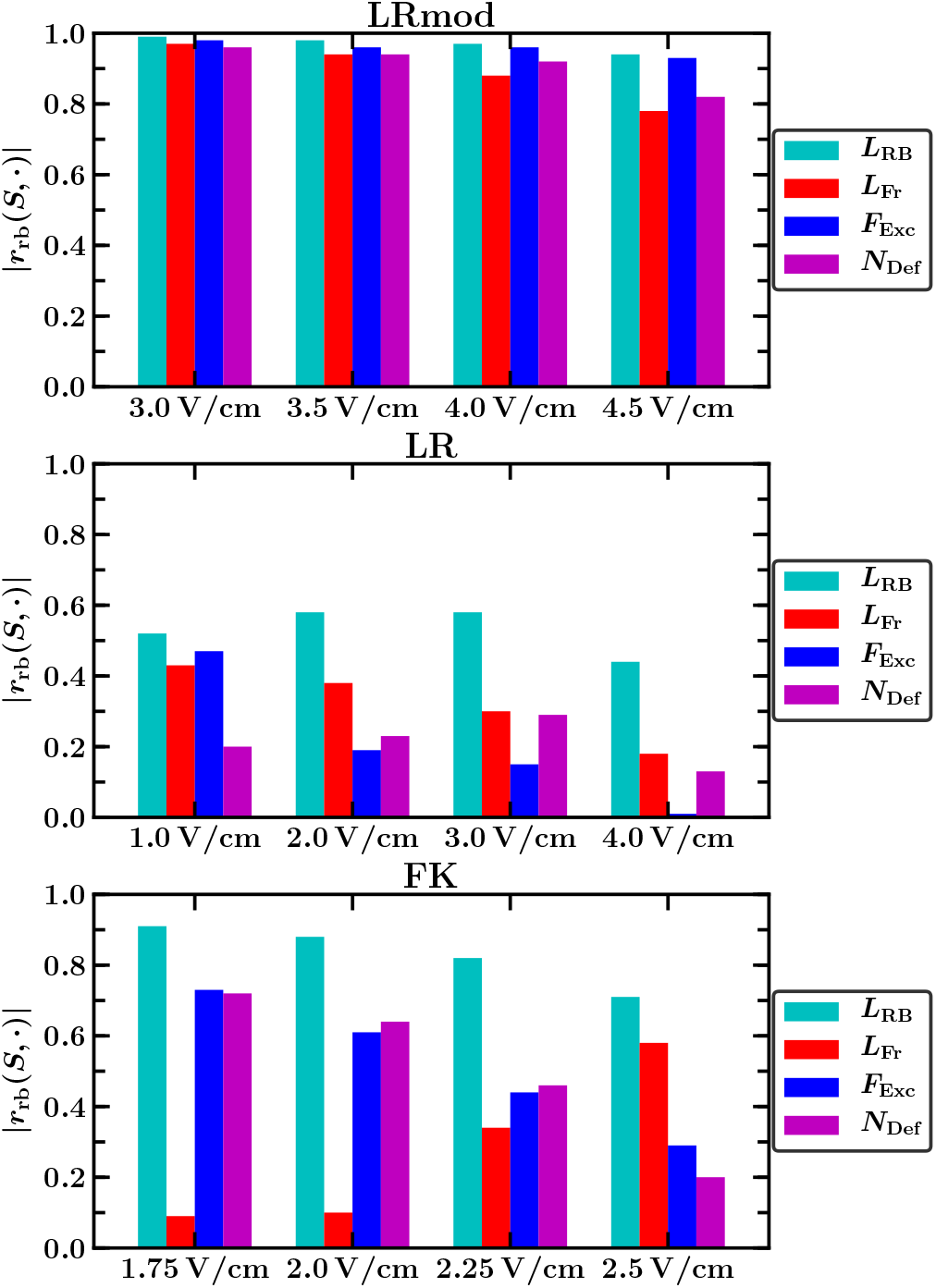
Absolute values of the *Rank Biserial Correlation Coefficients* |*r*_rb_| for the (LRmod), (LR) and (FK) model and different electric field strength *E* between defibrillation success *S* and the fraction of excitable *F*_Exc_, refractory boundary length *L*_RB_, front length *L*_Fr_ and number of defects *N*_Def_. The strongest correlations with the defibrillation success *S* were found for the refractory boundary length *L*_RB_. The refractory boundary length *L*_RB_ is furthermore the only macroobservable at all that is highly correlated with *S* for all three models and field strength *E* and therefore the macro-observable that can be used to predict the success probability for all three models.

**TABLE I.** *Rank Biserial Correlation Coefficients r*_rb_ for the (LRmod), (LR) and (FK) model and different electric field strength *E* between defibrillation success *S* and the following eleven macro-observables: fraction of excitable *F*_Exc_, refractory boundary length *L*_RB_, front length *L*_Fr_, time derivative of the fraction of excitable *∂*_*t*_*F*_Exc_, time derivative of the refractory boundary length *∂*_*t*_*L*_RB_, time derivative of the front length *∂*_*t*_*L*_Fr_, number of excitable cluster *N*_cExc_, number of non-excitable cluster *N*_cNExc_, number of refractory boundaries *N*_RB_, number of fronts *N*_Fr_ and number of defects *N*_Def_. The highest correlations are marked (**red**) while comparable high correlations are marked (**orange**). Most of the highest correlations were found for the refractory boundary length *L*_RB_ which is furthermore the only macro-observable at all that is highly correlated with the defibrillation success *S* for all three models and field strength *E*. Overall, the refractory boundary length *L*_RB_ is the most promising macro-observable that is suitable to predict the success of a low-energy pulse for all three models and might therefore be useful to decipher the underlying mechanism behind LEAP.

| $E$ [V/cm] | LRmod | | | | LR | | | | FK | | | |
| --- | --- | --- | --- | --- | --- | --- | --- | --- | --- | --- | --- | --- |
|  | 3.0 | 3.5 | 4.0 | 4.5 | 1.0 | 2.0 | 3.0 | 4.0 | 1.75 | 2.0 | 2.25 | 2.5 |
| $r_{rb}(S, F_{Exc})$ | **0.98** | **0.96** | **0.96** | **0.93** | **0.47** | 0.19 | 0.15 | 0.01 | **0.73** | 0.61 | 0.44 | 0.29 |
| $r_{rb}(S, L_{RB})$ | **-0.99** | **-0.98** | **-0.97** | **-0.94** | **-0.52** | **-0.58** | **-0.58** | **-0.44** | **-0.91** | **-0.88** | **-0.82** | **-0.71** |
| $r_{rb}(S, L_{Fr})$ | **-0.97** | **-0.94** | **-0.88** | **-0.78** | **0.43** | 0.38 | 0.30 | 0.18 | -0.09 | 0.10 | 0.34 | 0.58 |
| $r_{rb}(S, \partial_t F_{Exc})$ | 0.02 | -0.01 | -0.02 | -0.05 | -0.30 | **-0.43** | **-0.47** | **-0.41** | -0.52 | **-0.71** | **-0.74** | **-0.81** |
| $r_{rb}(S, \partial_t L_{RB})$ | -0.47 | -0.52 | -0.56 | -0.55 | -0.14 | -0.17 | -0.13 | -0.18 | 0.12 | 0.08 | 0.05 | 0.00 |
| $r_{rb}(S, \partial_t L_{Fr})$ | -0.22 | -0.23 | -0.21 | -0.24 | 0.05 | -0.09 | -0.10 | -0.15 | 0.00 | 0.22 | 0.51 | 0.56 |
| $r_{rb}(S, N_{cExc})$ | **-0.94** | **-0.93** | **-0.92** | **-0.89** | **-0.49** | -0.35 | -0.30 | -0.09 | 0.05 | 0.04 | 0.17 | 0.23 |
| $r_{rb}(S, N_{cNExc})$ | -0.78 | -0.78 | -0.72 | -0.68 | 0.01 | 0.00 | 0.00 | 0.00 | -0.20 | -0.24 | -0.19 | -0.20 |
| $r_{rb}(S, N_{RB})$ | **-0.98** | **-0.97** | **-0.96** | **-0.89** | **-0.44** | -0.38 | -0.35 | -0.13 | -0.66 | -0.59 | -0.44 | -0.22 |
| $r_{rb}(S, N_{Fr})$ | **-0.98** | **-0.96** | **-0.94** | **-0.84** | -0.16 | -0.18 | -0.25 | -0.08 | -0.67 | -0.58 | -0.39 | -0.12 |
| $r_{rb}(S, N_{Def})$ | **-0.96** | **-0.94** | **-0.92** | **-0.82** | -0.20 | -0.23 | -0.29 | -0.13 | **-0.72** | -0.64 | -0.46 | -0.20 |

**TABLE II.** *Spearman*’*s Rank Correlation Coefficient r*_s_ for the (LRmod), (LR) and (FK) model and different field strength *E* between refractory boundary length *L*_RB_ and fraction of excitable *F*_Exc_, front length *L*_Fr_, number of excitable clusters *N*_cExc_, number of refractory boundaries *N*_RB_, number of fronts *N*_Fr_ and number of defects *N*_Def_ as well as between time derivative of the fraction of excitable *∂*_*t*_*F*_Exc_ and difference between total length of the refractory boundaries and excitation fronts (*L*_RB_- *L*_Fr_). The six macro-observables in the upper part of the table (*F*_Exc_, *L*_Fr_, *N*_cExc_, *N*_RB_, *N*_Fr_ and *N*_Def_) are all relatively high correlated with *L*_RB_ for the (LRmod) model. *∂*_*t*_*F*_Exc_ in the lower part of the table is highly correlated with the difference (*L*_RB_-*L*_Fr_) for all three models.

| $E$ [V/cm] | LRmod | | | | LR | | | | FK | | | |
| --- | --- | --- | --- | --- | --- | --- | --- | --- | --- | --- | --- | --- |
|  | 3.0 | 3.5 | 4.0 | 4.5 | 1.0 | 2.0 | 3.0 | 4.0 | 1.75 | 2.0 | 2.25 | 2.5 |
| $r_s(F_{Exc}, L_{RB})$ | -0.75 | -0.74 | -0.73 | -0.72 | 0.00 | 0.02 | -0.01 | -0.04 | -0.61 | -0.57 | -0.55 | -0.55 |
| $r_s(L_{Fr}, L_{RB})$ | 0.54 | 0.53 | 0.50 | 0.45 | -0.31 | -0.16 | -0.11 | -0.08 | -0.29 | -0.35 | -0.34 | -0.33 |
| $r_s(N_{cExc}, L_{RB})$ | 0.71 | 0.73 | 0.76 | 0.77 | 0.45 | 0.46 | 0.45 | 0.43 | 0.14 | 0.11 | 0.06 | 0.03 |
| $r_s(N_{RB}, L_{RB})$ | 0.81 | 0.84 | 0.84 | 0.83 | 0.59 | 0.60 | 0.61 | 0.58 | 0.57 | 0.54 | 0.51 | 0.50 |
| $r_s(N_{Fr}, L_{RB})$ | 0.70 | 0.73 | 0.72 | 0.69 | 0.38 | 0.40 | 0.44 | 0.39 | 0.47 | 0.43 | 0.39 | 0.38 |
| $r_s(N_{Def}, L_{RB})$ | 0.71 | 0.75 | 0.74 | 0.73 | 0.42 | 0.43 | 0.47 | 0.42 | 0.52 | 0.48 | 0.44 | 0.43 |
| $r_s(\partial_t F_{Exc}, L_{RB} - L_{Fr})$ | 0.72 | 0.77 | 0.80 | 0.82 | 0.70 | 0.65 | 0.63 | 0.62 | 0.78 | 0.80 | 0.79 | 0.78 |

Beside *L*_RB_, there are further macro-observables that have comparably high correlation with *S* for at least one of the three considered cellular models. All of these correlations have in common that they are tightly linked with the refractory boundary length *L*_RB_. For example the high correlations of the defibrillation success *S* with the front length *L*_Fr_, the fraction of excitable *F*_Exc_ and the number of defects *N*_Def_ for the (LRmod) model, see Fig.4, is based on the fact, that the (LRmod) model exhibits stable spirals with a typical characteristic size, width and curvature where typical size relations among these three macro-observables and *L*_RB_ exist, see Fig.3. For more details and an analysis of all considered macro-observables, see Appendix B 2.

Overall, the refractory boundary length *L*_RB_ turns out to be clearly the most promising macro-observable to decipher the underlying mechanism behind LEAP as it is the only tested macro-observable that exhibits strong correlation with successful defibrillation for all three models, [it i. e. small values of *L*_RB_ indicate a high probability for defibrillation. In the next section, we will explore this qualitative observation in a more quantitative manner.

### B. Functional relationship between the refractory boundary length and the termination probability

In order to reveal the functional relationship between the refractory boundary length *L*_RB_ and the termination probability *P*, we first had to bin the data, since the termination probability *P* itself can only be determined as the fraction of successful termination events. For this purpose, we sorted for each model and electric field strength *E* all the 10000 performed simulations runs by *L*_RB_, divided them into equally sized bins with a size of 200 and computed for each bin the termination probability *P* and the mean of the refractory boundary length *L*_RB_. As shown in Fig.5, the termination probability *P* decays for all three models and field strengths *E* exponentially with the refractory boundary length *L*_RB_ and a with the field strength *E* decreasing decay rate *k* (*E*):

**FIG. 5.**
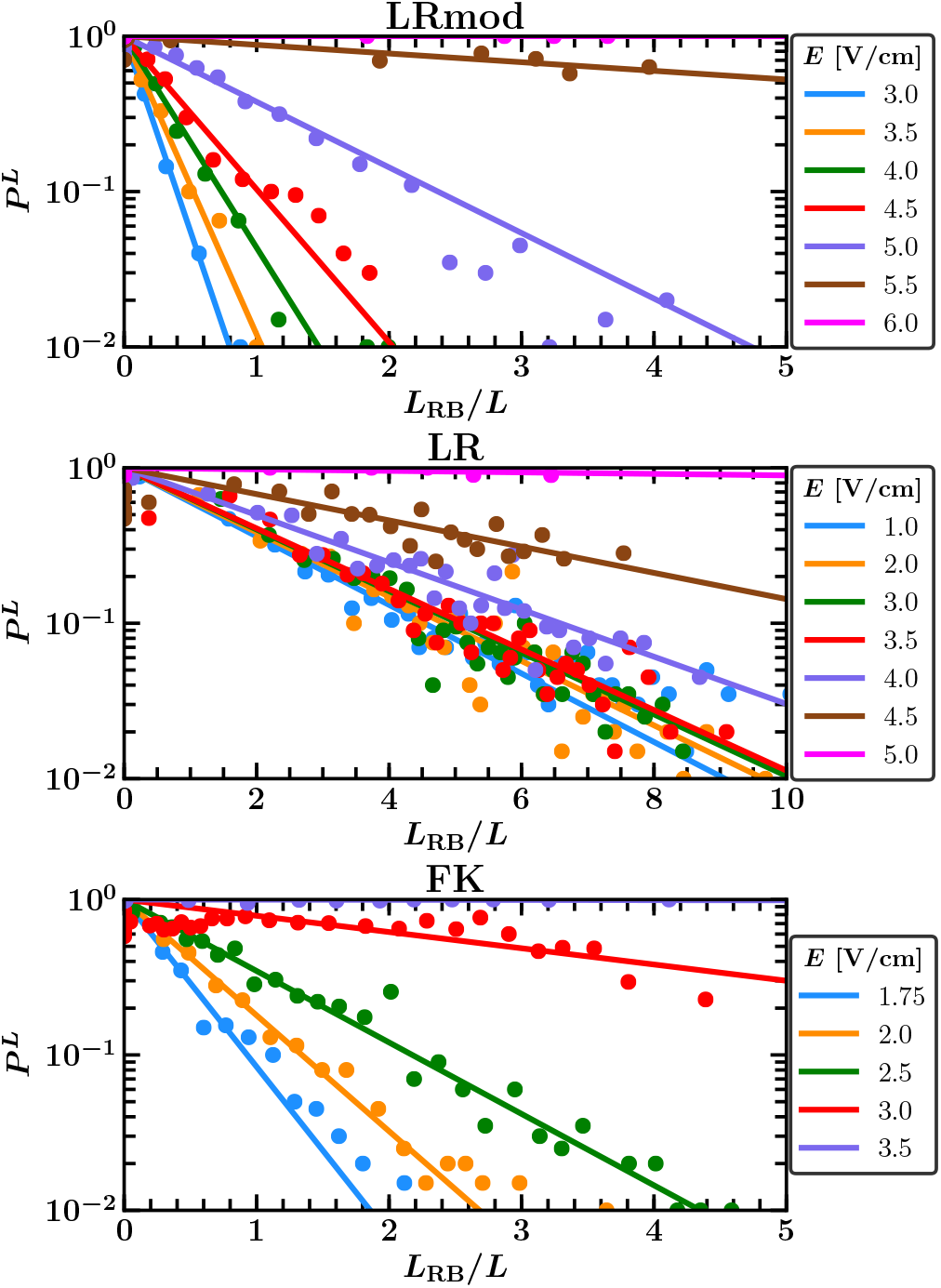
Scatter plot between the termination probability *P* and the normalized refractory border length *L*_RB_*/L* right before the pulse for different field strength *E. P* decays for each of the three models exponentially with *L*_RB_ and a with the field strength *E* decreasing decay rate. The exponential regressions of the data points for each field strength *E* are plotted as solid lines in the corresponding color.

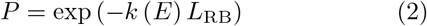

This decreasing decay rate *k* (*E*) becomes zero for field strength *E* greater than the single pulse defibrillation threshold and adds the field dependence. It reflects already that the termination probability *P* increases usually with the field strength *E* and that a pulse is beyond that always successful (*P* = 1) for field strength *E* greater than the single pulse defibrillation threshold, independent of the underlying dynamical state.

### C. Understanding the exponential relation between the refractory boundary length and the termination probability

We know now that the functional relationship between the termination probability *P* and the refractory boundary length *L*_RB_ is exponential, but not why and how we can interpret the field dependent decay rate *k* (*E*). In order to answer this question, we will first take a closer look on what is going on right after a pulse during successful LEAP, especially at the refractory boundaries, and derive then the exponential relation. The mechanisms, that determine the success of a pulse, are similar for all three models, albeit they are somewhat more involved for the (LR) and (FK) model. We will therefore first focus on the (LRmod) model and discuss the differences to the (LR) and (FK) model afterwards.

#### 1. A close look behind the scenes at the refractory boundaries during successful LEAP

Fig.6 shows for the (LRmod) model a sequence of snapshots right after the third pulse of a successful LEAP protocol with five pulses. Starting from the state right before the pulse (Fig.6a), we observe first the induction of excitation waves at the big heterogeneities (Fig.6b), which excite quickly the entire excitable tissue (Fig.6c) until almost all excitation waves die out (Fig.6d). All previous existing excitation fronts are terminated and only a few of the new generated excitation fronts, that follow the refractory boundaries into the progressively recovering parts of the tissue, survive (Fig.6e). These newly induced excitation fronts grow and increasingly roll in and form the new spirals (Fig.6f-h). Thus, each LEAP pulse terminates actually all previous existing excitation fronts, however, induces new excitation fronts at the refractory boundaries as well.

**FIG. 6.**
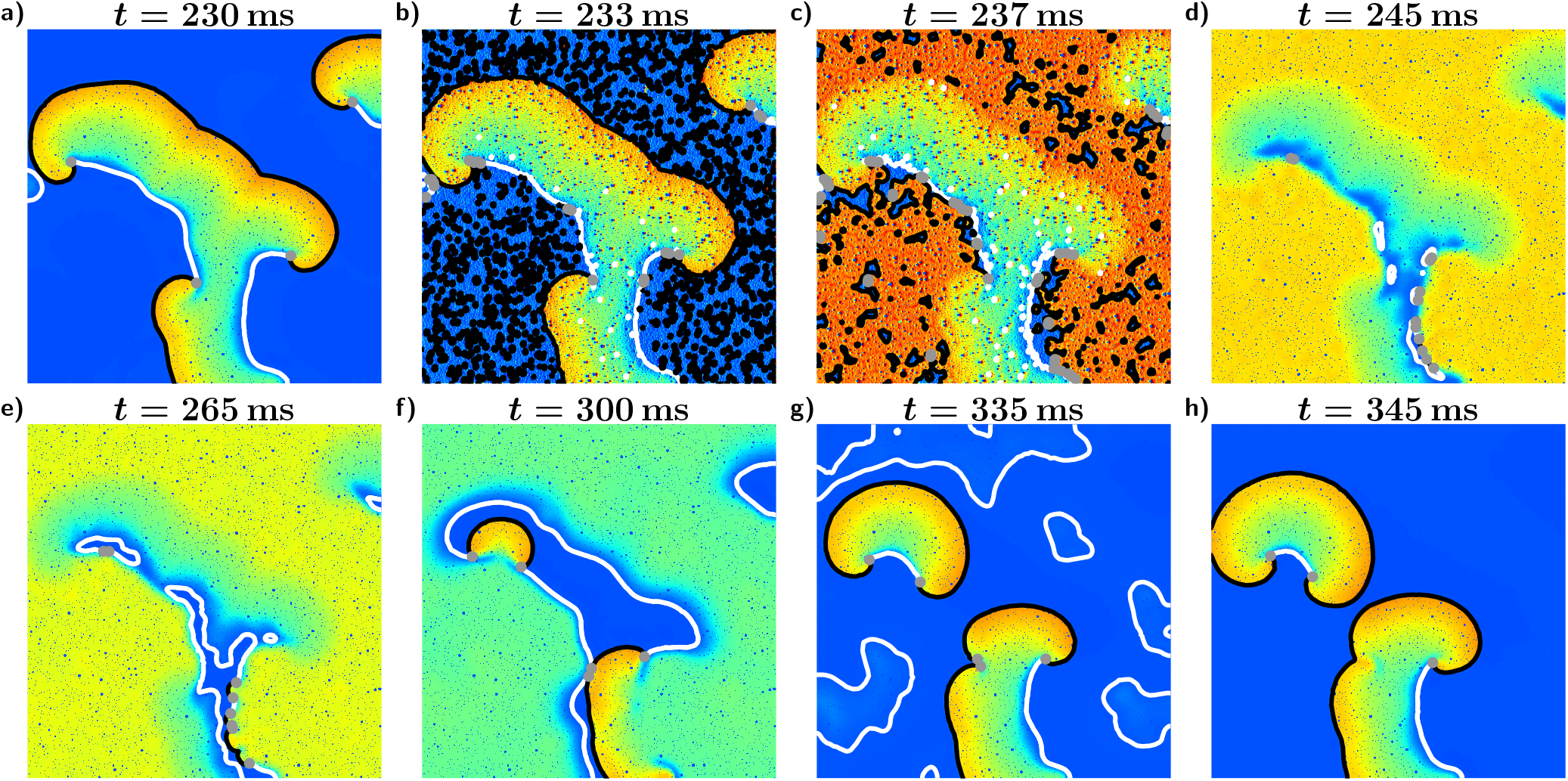
Sequence of snapshots of the transmembrane potential for a state of the (LRmod) model right after the third pulse of a LEAP protocol with a pacing period of *T* = 115 ms and field strength *E* = 3.5 V*/*cm, see also provided videos in the supplementary material. Excitation fronts are plotted black, refractory boundaries are plotted white and the topological defects are marked as grey dots. (a) State right before the pulse. (b) The pulse induces excitation fronts by depolarizing the tissue strong enough at the big heterogeneities. (c) The induced fronts excite quickly the entire excitable tissue. (d) Almost all excitation fronts die out. (e-f) All previous existing excitation fronts are terminated and only a few of the new generated excitation fronts, that follow the refractory boundaries into the progressively recovering parts of the tissue, survive. (g) These newly induced excitation fronts grow and increasingly roll in and form the new spirals. The refractory parts fall apart into smaller pieces and their corresponding margins shrink faster than the backs of the new spirals grow. (h) State right before the next pulse.

The mechanism that is relevant for the induction of these new excitation fronts is illustrated exemplary for a plane wave in Fig.7. Right after a pulse, the electric field induces first, as already mentioned, excitation fronts at the sufficient big heterogeneities (Fig.7b), that annihilate quickly each other and all previous existing excitation fronts (Fig.7c). Almost the entire tissue is now excited and only fronts, following the refractory boundaries into the progressively recovering parts of the tissue might survive. On or at least close to the refractory boundary, however, there are heterogeneities, that induce excitation fronts, which can only spread away from the refractory boundary and block therefore all fronts that move towards the refractory boundary at these positions. Thus, the newly induced excitation fronts are given by those fronts, that can break trough at places, where the refractory boundary is not sufficiently covered by activation sites that induce these blocking excitation fronts (Fig.7d). The heterogeneities and therefore those activated tissue sites as well are evenly distributed and the expected total number of newly induced excitation fronts right after a pulse is thus proportional to the total length of all refractory boundaries *L*_RB_:

**FIG. 7.**
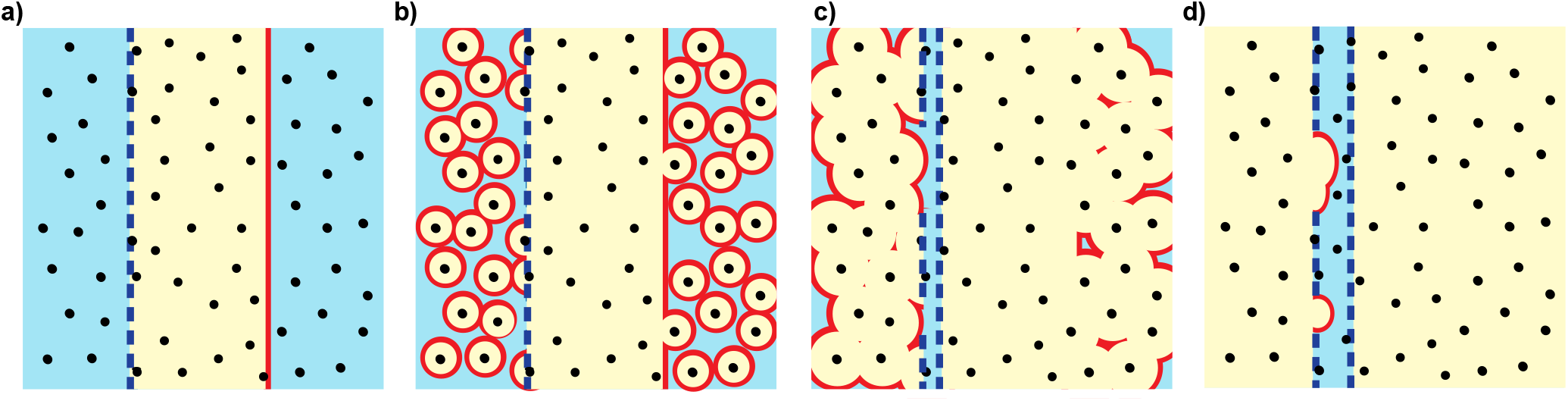
Illustration of the process, that affects right after a pulse the induction of new excitation fronts at the refractory boundaries, exemplary for a plane wave. Excitable tissue is colored blue, non-excitable tissue is colored yellow, excitation fronts are plotted red, refractory boundaries are plotted dashed blue and heterogeneities are marked black. (a) State right before the pulse. (b) The pulse induces excitation fronts at the big heterogeneities. (c) The induced fronts annihilate quickly each other and the entire front of the plane wave. Only fronts moving towards the refractory boundary might survive. On or at least close to the refractory boundary, however, there are heterogeneities, that have induced excitation fronts, which can only spread away from the refractory boundary and therefore block all fronts that move towards the refractory boundary at these positions. (d) Almost the entire tissue is now excited. The newly induced excitation fronts are given by those fronts, that can break trough at places, where the refractory boundary is not sufficiently covered by activation sites that induce these blocking excitation fronts.

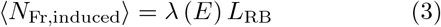

The proportionality factor *λ* (*E*) is the line density of newly induced excitation fronts along the refractory boundaries and usually decreases with the field strengt *E*, as can be seen in Appendix C exemplary for the back of a plane wave. That is because the minimal radius, necessary to activate a heterogeneity decreases for high field strength *E*.^27–29^ This leads to an increase of the density of activated heterogeneities, that cover the refractory boundaries, and thus to a reduction of excitation fronts that can break through. *λ* (*E*) should further become zero for field strength *E* greater than the single pulse defibrillation threshold, since a pulse is for this field strengths always successful and therefore, the expected total number of induced excitation fronts ⟨*N*_Fr,induced_⟩ has to be zero, independent of the current refractory boundary length *L*_RB_. Overall, the line density *λ* (*E*) features about all in Sec III B stated properties of the decay rate *k* (*E*) and as we will see in the following, substantially determines the decay rate *k* (*E*).

#### 2. Derivation of the exponential relation between the refractory boundary length and the termination probability

We found, that each LEAP pulse terminates all previous existing excitation fronts, however, induces new excitation fronts at the refractory boundaries as well, which arouse new fibrillation. Thus, the termination probability of a pulse *P* can be interpreted as the probability that no new excitation front will be induced. In order to determine this probability, we will first divide all refractory boundaries into *n* elements of equal length *L*_RB_*/n*. Analogous to the expected total number of newly induced excitation fronts ⟨*N*_Fr,induced_⟩ in equation (3), the expected total number of newly induced excitation fronts at each individual element is given by *λ* (*E*) *L*_RB_*/n*, which can be interpreted, if *n* is large enough and therefore this quotient is ≤1, as the probability at each individual element that a new excitation front will be induced. The probability at each individual element that no new excitation front will be induced is then simply given by the complementary probability 1 − *λ* (*E*) *L*_RB_*/n* and consequently the probability that no new excitation front will be induced at any of these elements is given by:

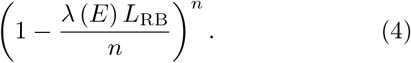

The termination probability *P* or rather the probability, that no new excitation front will be induced, is then given by the continuous limit *n* → ∞, which finally leads to the looked for exponential relation with the refractory boundary length *L*_RB_:

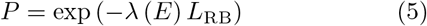

Hence, comparing with equation (2), the decay rate *k* (*E*) should be equal to the line density of newly induced excitation fronts *λ* (*E*). That is indeed the case as shown in Fig.8, where *k* (*E*) were determined as the decay rate of the exponential regression between the termination probability *P* and the refractory boundary length *L*_RB_ and *λ* (*E*) were determined as the average ratio between the number of induced excitation fronts right after a pulse and the corresponding refractory boundary length right before the pulse, further details can be found in Appendix D. Comparing further equation (5) with equation (3), one can say, that the termination probability *P* decays exponentially with the expected total number of newly induced excitation fronts ⟨*N*_Fr,induced_⟩:

**FIG. 8.**
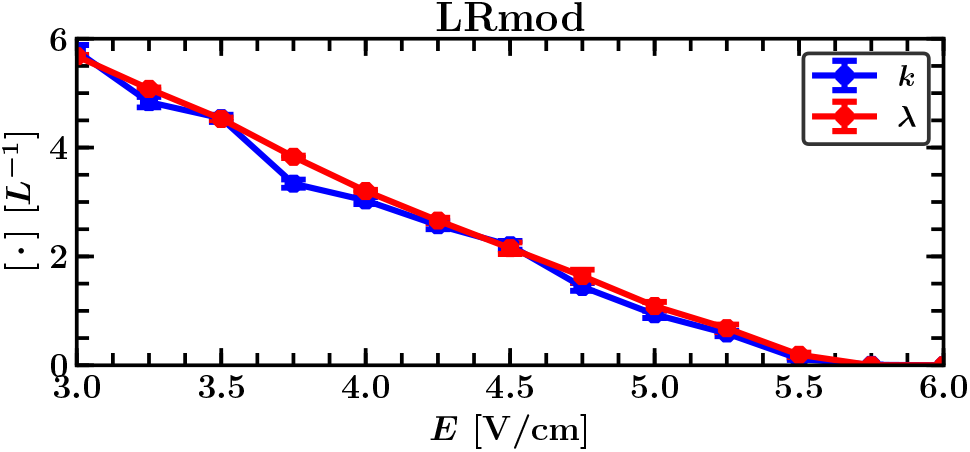
Decay rate of the exponential relation between the termination probability of a low-energy pulse and the refractory boundaray length right before the pulse *k* (blue) and line density of newly induced excitation fronts *λ* (red) as a function of the electric field strength *E* for the (LRmod) model. Both quantities are almost equal and decrease monotonically until they become zero for field strength *E* higher than the single defibrillation threshold of 6.0 V*/*cm.

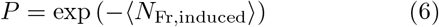

#### 3. Differences to the (LR) and (FK) model

So far we focused on the (LRmod) model and thus only on the defibrillation of stable spirals. Models that exhibit stable spirals have the particular feature, that excitation fronts usually do not run into refractory boundaries. Otherwise spirals could split up by hitting a refractory boundary or die out by moving into a dead end. The (LR) and the (FK) model, however, show a more complicated dynamic. They exhibit spatiotemporal chaos and unstable spirals, respectively, where excitation fronts, that run into refractory boundaries, are an integral part of the dynamic, see movies in the supplementary material, which has an additional significant impact on the defibrillation success, as we will show in the following.

Fig.9 shows for the (LR) model a sequence of snapshots right after the fourth pulse of a successful LEAP protocol with five pulses. Similar to the (LRmod) model, we observe here again, that the pulse terminates on the one hand all previous existing excitation fronts, however, induces on the other hand new fronts at the refractory boundaries which are given by those fronts that can follow a refractory boundary right after the pulse into the progressively recovering parts of the tissue. Besides getting blocked by excitation fronts that spread away from the refractory boundary, there exist a second process within the (LR) model, that significantly affects whether those fronts can follow. Refractory boundaries can move so slowly such that a following excitation front dies out by simply running directly into the refractory boundary itself, see Fig.9b-c (red encircled fronts). Whether a refractory boundary moves quick enough or not should not depend on the strength of the applied electric field. This second process is therefore field independent and requires only, that the pulse is strong enough to excite the entire excitable tissue. It dominates therefore for low field strength *E* the whole formation process of the newly induced excitation fronts, since the refractory boundaries are not sufficiently covered by activated heterogeneities for this low field strengths *E* anyway to completely block any of the following excitation fronts. Compared to the (LRmod) model, the line density of newly induced excitation fronts *λ* is thus for the (LR) model almost constant within the range of low field strength *E*, see Fig.10a. At first a field strength of *E* = 3.75 V*/*cm it starts to behave like for the (LRmod) model and decreases monotonically until it becomes zero for field strength *E* greater than the single defibrillation threshold.

**FIG. 9.**
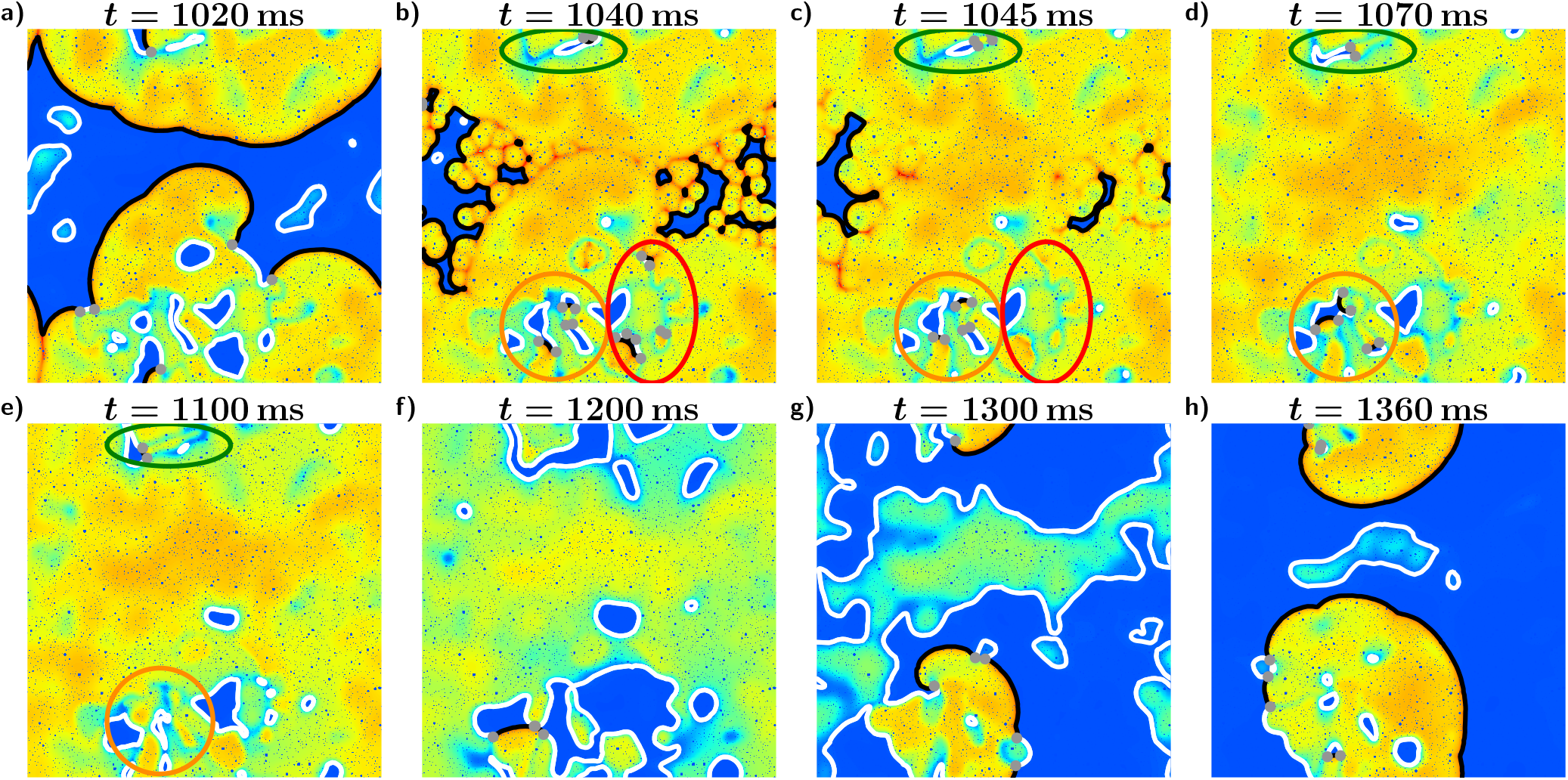
Sequence of snapshots of the transmembrane potential for a state of the (LR) model right after the fourth pulse of a LEAP protocol with pacing period *T* = 340 ms and field strength *E* = 1.0 V*/*cm, see also provided videos in the supplementary material. Excitation fronts are plotted black, refractory boundaries white and topological defects are marked as grey dots. (a) State right before the pulse. (b-c) The pulse induces excitation fronts at the big heterogeneities, which excite quickly the entire excitable tissue, until almost all excitation fronts die out. Only fronts moving towards a refractory boundary might survive, depending on whether the tissue along the refractory boundary is recovering quick enough and the front can move into the progressively recovering parts of the tissue (green or orange), or not and the front ends at a dead end (red). (d-e) These newly induced fronts can later on still disappear, annihilating each other or ending at a dead end (orange). In this example only one of the newly induced excitation fronts survived (green). (f-g) This front creates new spatiotemporal chaos. (h) State right before the next pulse.

**FIG. 10.**
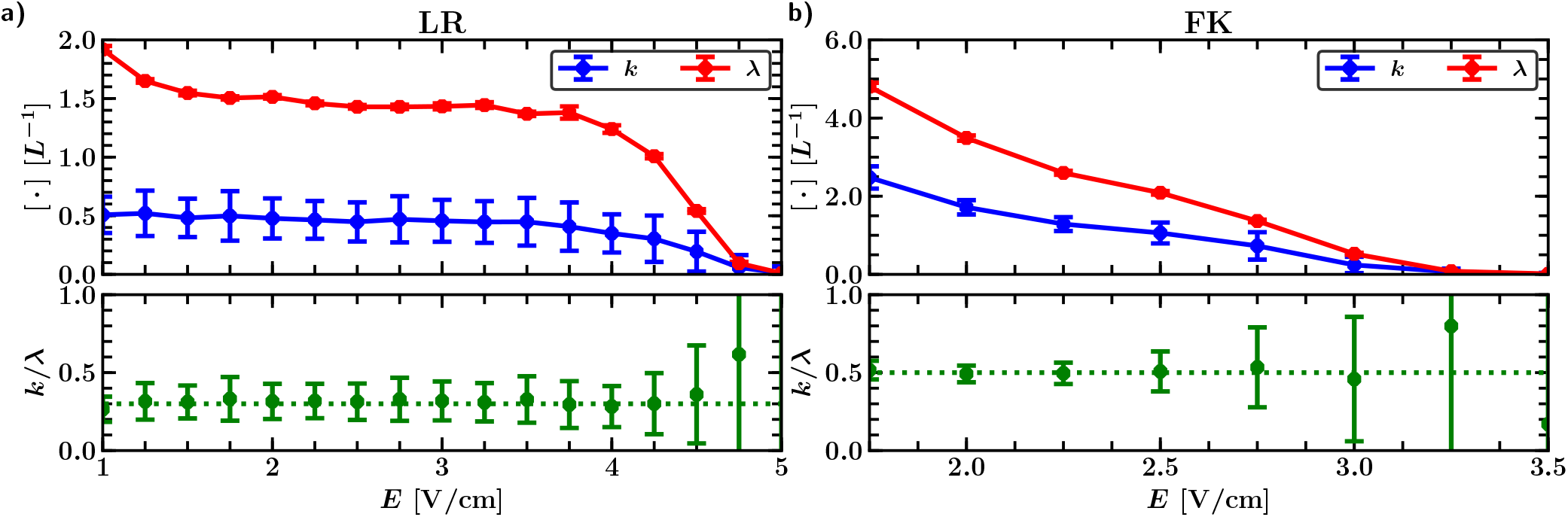
Decay rate *k* of the exponential relation between the termination probability of a low-energy pulse and the refractory boundary length right before the pulse (blue) and line density of newly induced excitation fronts *λ* (red) as a function of the electric field strength *E* for (a) the (LR) and (b) the (FK) model. Both, *k* and *λ* decrease for the (FK) model, just like for the (LRmod) model (Fig.8), monotonically until they become zero for field strength *E* higher than the single defibrillation threshold, whereas for the (LR) model, those two quantities remain first almost constant for field strengths *E* below 3.75 V*/*cm and start then to monotonically decrease until they become zero for field strength *E* higher than the single defibrillation threshold. Both, the (FK) and (LR) model have in common that *k* and *λ* are, compared to the (LRmod) model (Fig.8), not equal but rather proportional. The quotient of *k* and *λ* (green) is roughly constant with a value of 0.3 for the (LR) and 0.5 for the (FK) model.

For the (FK) model, this second process of refractory boundaries, that move so slowly such that following excitation fronts die out, is not observable. The line density of newly induced excitation fronts *λ* decreases therefore just like for the (LRmod) model monotonically until it becomes zero for field strength *E* greater than the single defibrillation threshold, as shown in Fig.10b.

Besides, both, the (FK) and (LR) model have in common, that *k* and *λ* are proportional, but not equal, which contradicts the in Sec III C 2 made implication, that both quantities are equal. This inconsistency can be explained by the fact, that we have so far implicitly assumed that each newly induced excitation front would survive. Survive means in this context for a front, that this front or at least one in context with this front newly evolved successor front is part of the new fibrillation. This assumption was valid for the (LRmod) model, where all newly induced excitation fronts grow and fuse together to stable spirals, but does not hold for the (LR) and (FK) model, where newly induced excitation fronts can annihilate each other or die out in dead ends, as shown exemplary for the (LR) model in Fig.9d-e (orange encircled fronts). Sufficient for the defibrillation success is therefore already that none of the newly induced excitation fronts survive. Hence, we have to consider instead of all newly induced excitation fronts actually only those that survive. This can be done by replacing the expected total number of newly induced excitation fronts ⟨*N*_Fr,induced_⟩ by the one of those fronts that survive:

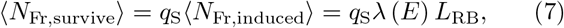

where *q*_S_ is the average survival rate of the newly induced excitation fronts. Doing so in equation 6, we get then, that the termination probability *P* decays actually exponentially with the expected total number of newly induced excitation fronts that survive *(N*_Fr,survive_):

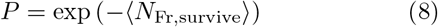

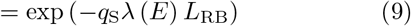

Comparing with equation (2), we get then that the decay rate *k* and the line density of newly induced excitation fronts *λ* are indeed proportional and that the proportionality factor should be given by the survival rate *q*_S_:

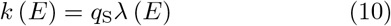

A verification whether the survival rate *q*_S_ is really the proportionality factor is hard, since a direct measurement of *q*_S_ would therefore be necessary, including the tracking of each newly induced excitation front and all their successor fronts. However, one can check whether the proportionality factor between *k* and *λ*, calculated as the quotient of *k* and *λ*, fulfills at least the properties that one would expect for a survival rate *q*_S_. This quotient of *k* and *λ* should then be field-independent, since the induction of the newly induced excitation fronts is usually completed after the application of the electric field. That is the case, as shown in Fig.10, the quotient of *k* and *λ* has roughly a constant value of 0.3 for the (LR) and 0.5 for the (FK) model. Apart from that, a lower value of the quotient of *k* and *λ* for the (LR) than for the (FK) model corresponds further with the observation, that the dying out of newly induced excitation fronts happens more frequently for the (LR) than for the (FK) model, see videos in the supplementary material.

### D. Understanding LEAP

So far we found, that the termination probability *P* decays exponentially with the expected total number of newly induced excitation fronts that survive ⟨*N*_Fr,survive_⟩. Furthermore, the survival rate of this newly induced excitation fronts is constant. Maximizing the termination probability of a pulse is therefore equivalent with minimizing the expected total number of newly induced excitation fronts ⟨*N*_Fr,induced_⟩ = *λ* (*E*) *L*_RB_. The easiest way to achieve this, is the usage of a field strength *E*, that is high enough, such that the line density of newly induced excitation fronts *λ* (*E*) is zero and thus no new excitation fronts get induced, independent of the underlying dynamical state or refractory boundary length *L*_RB_. That actually explains how single pulse defibrillation works and why the single pulse defibrillation threshold equals the field strengths where *λ* (*E*) starts to become zero. LEAP, however, operates in the range of much smaller field strength *E*, where *λ* (*E*) is significantly higher than zero and where considerable high termination probabilities *P* can therefore only be achieved for sufficient low refractory boundary lengths *L*_RB_. Hence, one pulse is not enough anymore and instead a coordinated interplay between several pulses is needed, that gradually decreases *L*_RB_ to values where defibrillation becomes likely. As shown in Fig.11, successful LEAP protocols achieve this coordinated interplay, by pulsing, when the refractory boundary length *L*_RB_ is minimal. This maximizes on the one hand the termination probability of each individual pulse, but more importantly, as we will show in the following, also maximizes the gradual decrease of *L*_RB_, and leads therefore overall to optimal defibrillation results.

**FIG. 11.**
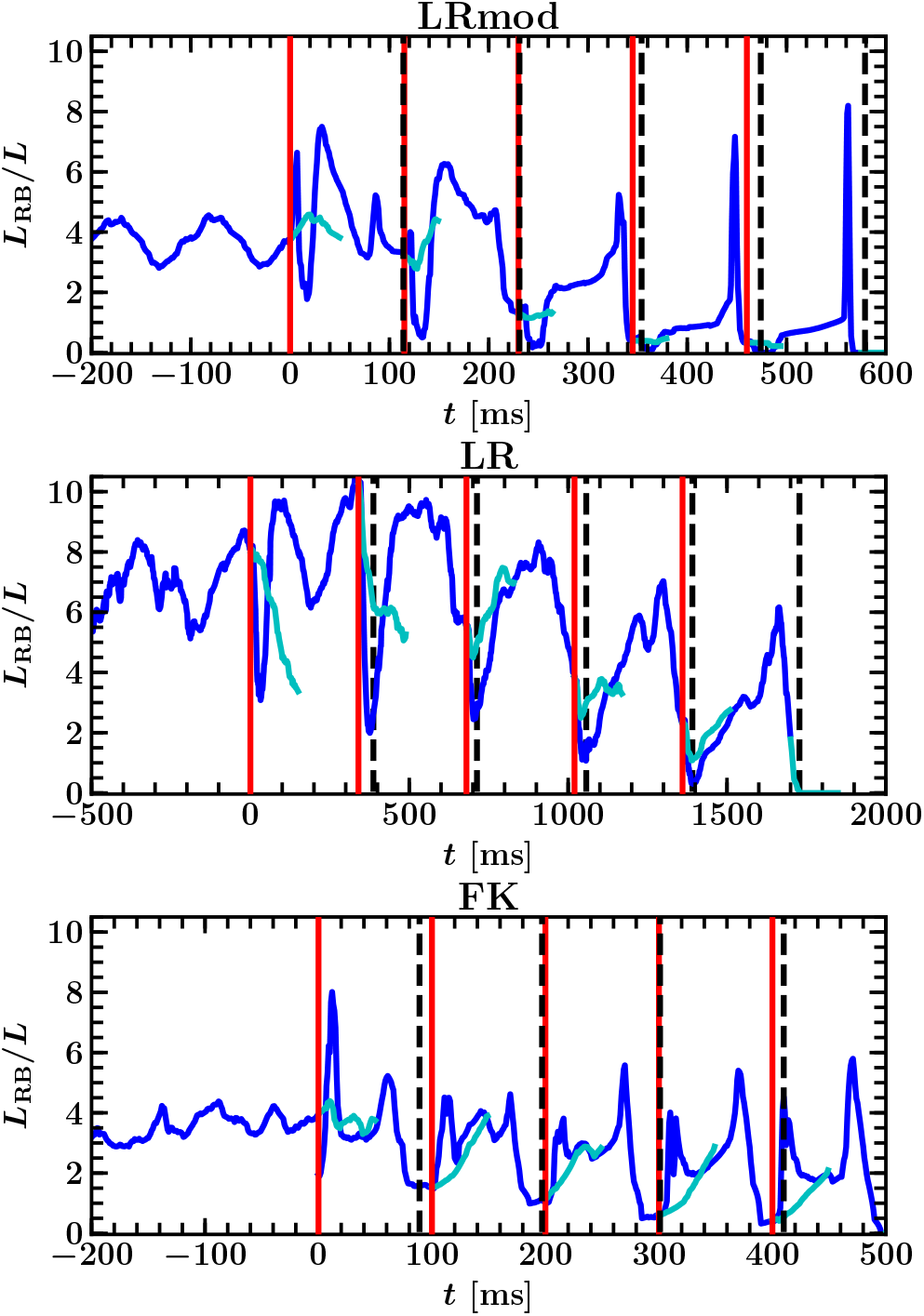
Development of the refractory boundary length *L*_RB_ (blue) over time *t* during successful LEAP with 5 pulses. The vertical red lines mark the times at which the pulses were applied, while in cyan is plotted how *L*_RB_ would evolve afterwards without the pulse. The shown successful LEAP episodes have for all three models in common, that the pulses roughly appear when *L*_RB_ and therefore the expected total number of newly induced excitation fronts is minimal which causes a gradual decrease of *L*_RB_ that simultaneously increases the success probability until complete defibrillation. Those minima of *L*_RB_ appear approximately when only refractory boundaries are left that are located at the backs of the newly induced excitation fronts (black vertical dashed lines). The applied LEAP protocols had a field strength and pacing period of *E* = 3.5 mV*/*cm and *T* = 115 ms for the (LRmod) model, *E* = 1.0 mV*/*cm and *T* = 340 ms for the (LR) model and *E* = 2.0 mV*/*cm and *T* = 90 ms for the (FK) model. Movies of the simulations are provided in the supplementary material.

#### 1. How the timing of the pulses affects the gradual decrease of the refractory boundary length during LEAP

We will now take a closer look on how the refractory boundaries evolve after a pulse during successful LEAP, in order to get a deeper insight into how the timing of the pulses affects the gradual decrease of *L*_RB_ during LEAP and especially why pulsing at minima of *L*_RB_ maximizes this gradual decrease of *L*_RB_. As shown in Fig.11, the refractory boundary length *L*_RB_ often reaches its lowest values shortly after the application of a pulse, when almost the entire tissue is excited and only the newly induced excitation fronts are left, which are at this stage still small and just started to grow, see exemplary Fig.6d for the (LRmod) or Fig.9d for the (LR) model. A pulse at this time, however, would be usually not very effective, since the remaining refractory boundaries have hardly moved and are therefore effectively still at those positions in the tissue located where the induction of the newly induced excitation fronts took place and thus a re-induction of those fronts would not be very unlikely. Thereafter the tissue starts increasingly to recover and the refractory boundaries become larger, see exemplary Fig.6e-g for the (LRmod) or Fig.9e-g for the (LR) model. The refractory boundary length *L*_RB_ reaches its maximal values when these refractory parts of the tissue start to split up into smaller pieces, whose refractory boundaries shrink after that faster than refractory boundaries that have started to emerge at the backs of the newly induced excitation fronts or their successor fronts grow, see exemplary Fig.6g-h for the (LRmod) or Fig.9g-h for the (LR) model. Hence, the minimum of *L*_RB_ after a pulse appears at the time when only refractory boundaries are left that are located at one of those backs (black dashed lines in Fig.11). This time when *L*_RB_ is minimal marks also the re-fibrillation threshold *T*_rFib_, since after a successful pulse, when the number of newly induced excitation fronts is zero, those backs would not exist and therefore, there would be exactly after this time no refractory boundaries left at which fibrillation could get re-induced. After the re-fibrillation threshold *T*_rFib_, the refractory boundary length *L*_RB_ is roughly proportional to the total number of newly induced excitation fronts *N*_Fr,induced_, since *L*_RB_ is then solely given by the total length of the refractory boundaries located at the backs of all newly induced excitation fronts and their successor fronts. This means for two arbitrary subsequent pulses during successful LEAP with a temporal spacing *T* ≥ *T*_rFib_, that the expected refractory boundary length right before the second of the two subsequent pulses 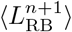 is proportional to the expected total number of newly induced excitation fronts 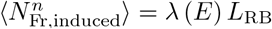 that get induced for a refractory boundary length *L*_RB_ that is equal to the length right before the first of the two subsequent pulses 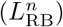:

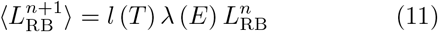

The expected ratio of the refractory boundary lengths of two subsequent pulses is then given by:

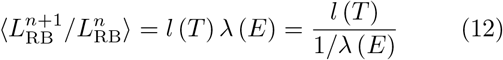

The proportionality factor *l* (*T*) can be interpreted as the average total length of the refractory boundaries after this temporal spacing *T* that were caused by a single newly induced excitation front. This length *l* is minimal for the re-fibrillation threshold *T*_rFib_ and depends actually only solely on *T* as long as the refractory boundaries grow independently of each other or more specifically as long as the newly induced excitation fronts have not started to fuse before exciting the tissue that forms after this spacing *T* the new refractory boundaries. That is the case as long as *l* is still significantly shorter than the average distance between the newly induced excitation fronts (roughly given by 1*/λ*, the reciprocal of the line density of newly induced excitation fronts) and thus satisfied by ratios 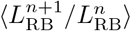 significantly smaller than 1, as can be seen in equation (12). Hence, for LEAP protocols that gradually decrease the refractory boundary length *L*_RB_ with temporal spacing *T* ≥ *T*_rFib_, those ratios 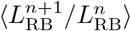 depend only on the field strength *E* and the current temporal spacing *T* itself and are therefore independent of the timing of the previous pulses. Consequently, a LEAP protocol that best possibly hit the minima of *L*_RB_ at the re-fibrillation threshold *T*_rFib_ minimizes all ratios 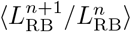, since *l* (*T*) is as already mentioned then minimal. That is basically why pulsing at minima of *L*_RB_ maximizes the gradual decrease of *L*_RB_ during successful LEAP.

#### 2. The range of successful LEAP protocols

Mandatory for the success of a LEAP protocol is, as already mentioned, a gradual decrease of *L*_RB_ to values where successful defibrillation becomes likely. Thus, successful LEAP protocols should be equal to those protocols, where the ratios 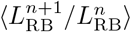 are significantly smaller than 1. The pacing periods where those ratios 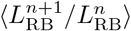 are minimal and consequently also the re-fibrillation threshold are roughly given by *T*_rFib_ = 115 ms for the (LRmod), *T*_rFib_ = 340 ms for the (LR) and *T*_rFib_ = 95 ms for the (FK) model, see Fig.12. Those re-fibrillation thresholds and the corresponding minimal field strengths, where 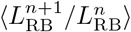 starts to become significantly smaller than 1 (vertical dotted lines), coincide indeed, as one would expect, with the pacing periods *T* and field strengths *E*, where LEAP starts to become successful, compare with Fig.2.

**FIG. 12.**
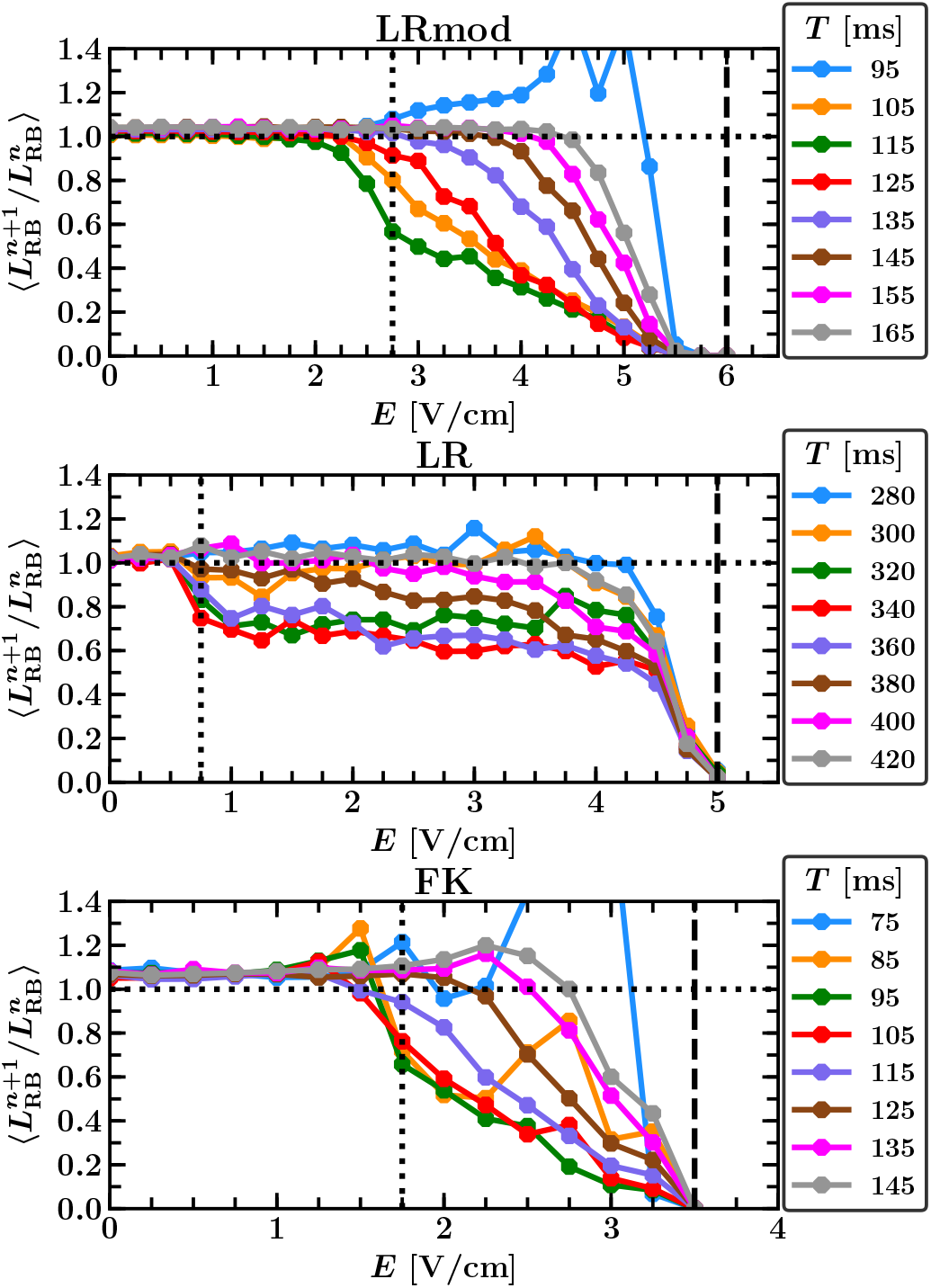
Average ratio 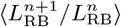 of the refractory boundary length *L*_RB_ between subsequent LEAP pulses as a function of the field strength *E* for different pacing periods *T*. Mandatory for the success of LEAP is a gradual decrease of *L*_RB_ and therefore ratios 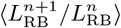 significantly smaller than 1 (horizontal dotted line). For weak field strength *E* all ratios have a value around one, since pulsing with such a weak field strength has nearly no impact. The minimal field strengths *E* where those ratios 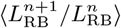 starts to become significantly smaller than 1 (vertical dotted lines) and the field strength *E* where all those ratios start to be zero independent of the used pacing periods *T* (vertical dashed lines) coincide with the minimal field strength *E* where LEAP starts to become successful and the threshold for single pulse defibrillation, respectively, compare with Fig.2. The re-fibrillation threshold, the pacing period *T* where those ratios 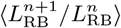 are minimal, is roughly given by *T*_rFib_ = 115 ms for the (LR-mod), *T*_rFib_ = 340 ms for the (LR) and *T*_rFib_ = 95 ms for the (FK) model. The range of pacing periods *T > T*_rFib_ with 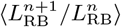 significantly smaller than 1 grows with increasing field strength *E* faster than the one for *T < T*_rFib_, reflecting that the margins of the refractory parts of the tissue increase faster towards lower pacing periods *T* than the backs of the newly induced excitation fronts grow for higher pacing periods *T*, exemplary shown in Fig.6g-h for the (LRmod) model. This explains why the range of successful pacing periods grows for *T > T*_rFib_ faster than for *T < T*_rFib_, see Fig.2.

For field strength *E* below these minimal field strengths (vertical dotted lines in Fig.12), all ratios 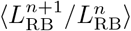 have a value around one, since pulsing with such a weak field strength has nearly no impact. Ratios 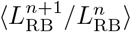 significantly smaller than 1 occur only for pacing periods *T* within a close range around the refibrillation threshold *T*_rFib_. For to high pacing periods *T* the ratios 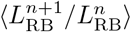 become one, see Fig.12, since the spacing between the pulses becomes then long enough for the newly induced excitation fronts to re-establish a new well-developed fibrillatory state after each pulse. To low pacing periods *T* on the other hand can lead partially even to ratios 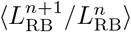 significantly higher than one, reflecting that the refractory boundary length *L*_RB_ consist then besides of the backs of the newly induced excitation fronts additionally on the margin of the fully refractory parts of the tissue that are not fully recovered after this temporal spacings *T < T*_rFib_.

The ratios 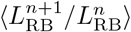 depend for LEAP protocols that successfully decrease *L*_RB_ with pacing periods *T* ≥ *T*_rFib_ only over the line density of newly induced excitation fronts *λ* (*E*) on the field strength *E*, as shown in the previous section (equation (12)). Thus, the ratios 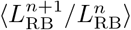 significantly smaller than 1 (horizontal dotted line in Fig.12) have for *T* ≥ *T*_rFib_ a similar course as *λ* (*E*), plotted in Fig.8 for the (LRmod) model and Fig.10 for the (LR) and (FK) model. That is why, just like *λ* (*E*), those ratios 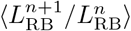 decrease for the (LRmod) and (FK) model monotonically with the field strength *E* until becoming zero for the single pulse defibrillation threshold (vertical dashed lines) and furthermore why those ratios 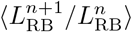 remain for the (LR) model first constant for field strengths *E* below 3.75 V*/*cm before they also start to decrease monotonically until becoming zero for the single defibrillation threshold. Overall, that explains for pacing periods *T* ≥ *T*_rFib_ why with an increasing field strength *E* the number of necessary pulses *n* for successful defibrillation decreases and why the range of pacing periods *T* with ratios 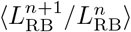 significantly lower than one and thus also the range of successful pacing periods *T* ≥ *T*_rFib_ increases until all pacing periods *T* start to be successful at the single pulse defibrillation threshold, compare Fig.12 with Fig.2.

Similar precise statements about the behaviour for pacing periods *T < T*_rFib_ are not possible, since the margins of the then not fully recovered fully refractory parts of the tissue play additionally an important role. One can only observe that the range of pacing periods *T < T*_rFib_ with 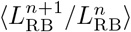 significantly smaller than 1 is usually smaller than the range for *T > T*_rFib_, see Fig.12. This reflects basically the already made observation from the previous section, that the margins of the refractory parts of the tissue become faster longer towards lower pacing periods *T* than the backs of the newly induced excitation fronts grow towards higher pacing periods *T*, exemplary shown in Fig.6g-h for the (LRmod) model, and explains why the range of successful pacing periods is for *T > T*_rFib_ bigger than for *T < T*_rFib_, see Fig.2.

## IV. DISCUSSION

The aim of the present study was to gain insight into the basic mechanisms behind defibrillation by periodic pacing with energies substantially lower than the single shock threshold energy for defibrillation. This approach was labelled as LEAP in earlier experimental studies^12,13^, a terminology which we have often adopted here to describe our simulation studies. Altogether, we have performed extensive simulations of two-dimensional models of a cardiac tissue perforated by randomly distributed heterogeneities of varying size representing blood vessels. The focus was on analysing the data with respect to the role of macro-variables typically for excitable media. Three different models with qualitatively different dynamics were used, namely, a modified Luo-Rudy model (LRmod) that exhibits stable spirals, the Luo-Rudy (LR) model and a version of the Fenton-Karma (FK) model that both exhibit phenomenologically different forms of spatiotemporal chaos. With the help of a large number of simulations, we were able to consider the potential relevance of a wide range of macro-observables for excitable media by determining their correlation with the success probability of an individual low-energy pulse during a LEAP pacing sequence. While our conclusion here are based on simulations in the relatively detailed 2D continuum model introduced in earlier publications^14,17^, related studies with a simplified continuum model^16,18^ and a cellular automaton^48,49^ show similar behavior in particular with respect to the occurence of an optimal pacing period equal to or slightly above the typical period of the fibrillatory dynamics pointing towards similar mechanism than the one at work in our model.

Our results in Sec III A and III B clearly show that the refractory boundary length *L*_RB_, the total length of the borders between refractory and excitable parts of the tissue, predict most reliably the success probability of a low-energy pulse during LEAP. The termination probability *P*_*L*_ of an individual low-energy pulse is found to depend exponentially on this length and a decay rate that decreases with the applied field strength, see Fig.5. *L*_RB_ was the only of the considered macro-observable that showed high correlation with defibrillation success for all three models. In contrast, we did not find convincing evidence that the number of defects *N*_Def_ is a good indicator for defibrillation as stipulated in alternative explanations that connect defibrillation with the unpinning and the removal of spirals from heterogeneities in the medium^11,15,27,34–36^. This does not imply that the number of defects is not important indicator for defibrillation. After all, fibrillation is of course driven by defects and defibrillation hence corresponds to removal of all defects. A small number of defects therefore increases the likelihood of successful defibrillation, since they are are more likely to get annihilated than a large number of defects. The number of defects, however, in contrast to the refractory boundary length cannot predict the probability for the emergence of new defects.

A closer look in Sec III C 1 revealed, that actually each pulse during LEAP terminates practically all the previously existing excitation fronts and defects. Pacing, however, often induced new rotor pairs and excitation fronts at the refractory boundaries. Almost all of these many small excitation fronts, that get generated right after a pulse at the sufficient large heterogeneities, again quickly annihilate each other. A few of these fronts on the other hand might nevertheless survive by trailing a refractory boundary into the recovering parts of the tissue, see Fig.6 and can develop into a new pair of topological defects. It is worth noting that a recent study by DeTal et al.^50^ showed that excitation along the entire contour of the refractory boundary is sufficient to terminate all excitation waves in the medium and thus provide successful defibrillation. In contrast, we observe that during pacing in the LEAP regime, some parts of the refractory boundary will not be excited immediately. In many cases these gaps are small and will rapidly closed by excitation waves from neighboring regions. Occasionally, however, these gaps are large enough to allow for the emergence of new pair of defects before the gap is closed. We found further, that the expected total number of newly induced excitation fronts is proportional to the refractory boundary length *L*_RB_ and a field dependent line density *λ* (*E*), which decreases with an increasing field strength *E*, since in this case the density of activated heterogeneities that block the incoming excitation fronts as well as the depolarization, that increase the recovering time of the surrounding tissue, becomes larger.

Overall, the termination probability *P* of an individual low-energy pulse can be interpreted as the probability that no new fibrillation arousing front gets induced. This leaded, as shown in Sec III C 2 and III C 3, to the observed exponential dependence between the success probability *P* and the refractory boundary length *L*_RB_ with a field dependent decay rate *k* (*E*) that is proportional to the line density of newly induced excitation fronts *λ* (*E*) as can be seen in Fig.8 for the (LRmod) and Fig.10 for the (LR) and (FK) model. The corresponding proportionality factor is interpretable as the ratio of newly induced excitation fronts that in average survive after a pulse and that are responsible for new fibrillation. This ratio was equal to one for the (LRmod) model, which exhibits stable spirals where all those newly induced excitation fronts survive, and it was below one for the (LR) and (FK) model, that have a dynamic, where some of those new fronts annihilate each other or simply die out by running into a refractory boundary instead of following it.

The interpretation of the termination probability as the probability that no new front gets induced is completely in conformity with the upper limit of vulnerability (ULV) hypothesis, which states for successful defibrillation, that a shock not only has to halt all excitation fronts that are present during fibrillation but also must not stimulate new excitation fronts that can re-fibrillate the heart.^51,52^ Several experimental studies^53–55^ have shown, that the so called upper limit of vulnerability, the highest field strength for a shock that is capable of initiating new fibrillation, does not differ significantly from the single pulse defibrillation threshold. This finding is reproduced in our simulations, where the defibrillation threshold of a single shock equals the field strength, where the line density of newly induced excitation fronts *λ* (*E*) becomes zero, see Fig.8 for the (LRmod) and Fig.10 for the (LR) and (FK) model. The measurement of the ULV limit usually takes place at a moment during the cardiac cycle where the initiation of new fibrillatory activity is most likely. This so called vulnerable window is located at the last half of the T wave when the ventricles have almost finished the repolarization. At this stage the fully refractory parts of the tissue disintegrate into smaller pieces and consequently the total length of the refractory boundaries at the corresponding margins reaches its maximal extent, as shown exemplary in Fig.6g-h for the (LRmod) model. Hence, the refractory boundary length and thus also the expected total number of newly induced excitation fronts become maximal during the vulnerable window. This explains why the heart is most vulnerable during this period of the cardiac cycle.

LEAP operates for field strengths substantially smaller than the ULV limit and for successful single-shock defibrillation and therefore requires a coordinated interplay between several pulses by gradually decreasing the refractory boundary length to values, where the probability of the induction of new excitation fronts becomes very small. The range of successful LEAP protocols is therefore roughly given by those protocols whose ratios between the refractory boundary length of subsequent pulses are much lower than one, as we have shown in Sec III D 2. We found further, as shown in Sec III D 1, that an optimal protocol requires pacing at times when the refractory boundary length is minimal, since this maximizes on the one hand the termination probability as well as the rate of decrease of the refractory boundary length. Those minima of the refractory boundary length appear roughly, as shown in Fig.11, at the time after a pulse at which the entire tissue, that got excited by the pulse, is completely recovered or more specifically at the time when the last refractory boundary that was located at the margin of one of the fully refractory parts of the tissue disappears. One can speculate that this recovering time should roughly coincide with the dominant period during fibrillation, since for a fully developed fibrillatory state the necessary time of the tissue to recover should be similar and tissue that becomes once again excitable should get very quickly re-excited during fibrillation as fibrillation is usually characterized by high frequencies. Taking further into account that the refractory boundaries located at the margins of those fully refractory parts of the tissue usually shrink faster than the refractory boundaries located at the backs of the newly induced excitation fronts grow, see exemplary Fig.6 for the (LR-mod) model, and that therefore the range of successful pacing periods usually increases with an increasing field strength faster towards higher pacing periods than towards lower pacing periods, as shown in Fig.2, explains why several experimental^12,13,32^ and numerical^14,17,32,33^ studies of LEAP provide evidence that the optimal observed pacing periods are in the range of the dominant arrhythmia cycle length or above. The presented results suggest that LEAP decreases gradually the refractory boundary length until successful defibrillation. Since there is a field dependent exponential relation between termination probability and refractory boundary length, is tempting to extend the picture to three dimensions (3D): the refractory boundary length would the become simply a refractory boundary surface and the line density of newly induced excitation fronts would change into a surface density, whereas defects or spirals would turn into filaments or scroll waves. The answer if the mechanism revealed here for two-dimensional (or quasi two-dimensional) excitable media like thin layers of excitable tissue will generalize into 3D will require, however, a much more extensive simulation study and is beyond the scope of our presented investigations. Another, important extension of our studies here would be the exploration of the role and impact of heterogeneities in the medium, e. g. representing fibrotic tissue or tissue damaged by infarctions, on the dynamics and success rates of defibrillation protocols.

The impression that spirals get eliminated one by one during successful LEAP and thus the number of defects, fronts or related quantities should play a crucial role for the success of a pulse, canbe explained simply by the fact that most but not all stable spirals get replaced after each pulse by excitation fronts that get induced at the backs of their spiral arms, see videos of the (LRmod) model in the supplementary material. Besides, the presented mechanism also rationalizes earlier interpretations of LEAP as a synchronization process, since a decrease of the refractory boundary length is often accompanied by an increase of the fraction of excitable tissue. Note, however that the picture presented by us is solely based on assumptions about local events and does not require involvement of a global ordering process as e. g. in the synchronisation of population of coupled oscillators. Overall, we are confident that the new insights presented in this work are crucial to understand the mechanism behind LEAP. The finding of the simple exponential relation between the defibrillation success rate and the length of the refractory boundary might help to develop better practical methods for a reliable estimation of a optimal pacing period or support the design of feedback controlled LEAP protocols.

## SUPPLEMENTARY MATERIAL

Movies can be obtained by request to the authors.

The simulation domain has in all movies no-flux boundary conditions for the (LRmod) model and periodic boundary conditions for the (LR) and (FK) model. In all movies, the (first) pulse is applied at *t* = 0 ms. All biphasic pulses are applied for 7 ms in the forward direction and 3 ms in the backward direction.

Movie 1: Defibrillation of a state of the (LRmod) model by LEAP with five biphasic pulses for a field strength *E* = 3.5 V*/*cm and pacing period *T* = 115 ms Movie 2: Defibrillation of a state of the (LRmod) model by LEAP with five biphasic pulses for a field strength *E* = 4.5 V*/*cm and pacing period *T* = 115 ms

Movie 3: Defibrillation of a state of the (LR) model by LEAP with five biphasic pulses for a field strength *E* = 1.0 V*/*cm and pacing period *T* = 340 ms

Movie 4: Defibrillation of a state of the (LR) model by LEAP with five biphasic pulses for a field strength *E* = 4.0 V*/*cm and pacing period *T* = 340 ms

Movie 5: Defibrillation of a state of the (FK) model by LEAP with five biphasic pulses for a field strength *E* = 2.0 V*/*cm and pacing period *T* = 100 ms

Movie 6: Defibrillation of a state of the (FK) model by LEAP with five biphasic pulses for a field strength *E* = 2.5 V*/*cm and pacing period *T* = 100 ms

Movie 7: Single pulse of a plane wave for the (LRmod) model and a field strength *E* = 3.0 V*/*cm

Movie 8: Single pulse of a plane wave for the (LRmod) model and a field strength *E* = 4.0 V*/*cm

Movie 9: Single pulse of a plane wave for the (LRmod) model and a field strength *E* = 5.0 V*/*cm

Movie 10: Single pulse of a plane wave for the (LRmod) model and a field strength *E* = 6.0 V*/*cm

## ACKNOWLEDGMENTS

We acknowledge financial support by DFG through SFB910 (PB, TN, MB) and through GRK1558 (PB).

## Appendix A: Analysis of discretization effects

As described in Sec II A, the heterogeneities in our composite model of two-dimensional homogeneous isotropic tissue represent non-conducting blood vessels that have circular shape. The size distribution of these circles is assumed to follow a power law, that was reported by Luther *et al*.^13^ for the vasculature of the heart muscle:

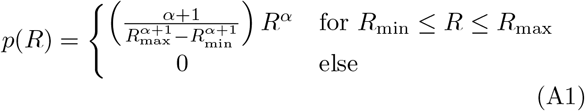

The density of activated heterogeneities for a given distribution of sizes (respectively radii) is obtained as:

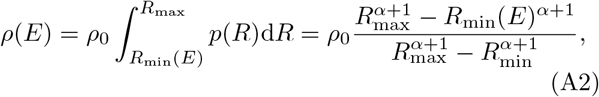

In all our simulations in the results section above, the smallest considered radius was *R*_min_ = 3 × 10^−4^*L*, the largest considered radius was *R*_max_ = 4 × 10^−3^*L*, the exponent was *α* = − 2.75 and the density of all heterogeneities was *ρ*_0_ = 1.6 × 10^4^, where *L* is the system length. The heterogeneity radii drawn from this distribution are close to the spatial resolution Δ*x* = 0.001 *L*, which was used for most of the simulations in this study. Heterogeneities with a diameter even smaller than the spatial resolution (2*R <* Δ*x*) were set to the size of one pixel. Such limitations in resolving circular heterogeneities are expected to mainly affect the activation behavior of the heterogeneities, since it may depend strongly on the size and shape of a heterogeneity used in the simulations^27–29^.

Resolution-related artifacts are clearly visible for the dependency of the minimal radius *R*_min_ (*E*) that is necessary for a heterogeneity to get activated by a pulse on the applied electric field strength *E*, see figure 13. The minimal activation radius *R*_min_ (*E*) decreases stepwise and not smoothly for the standard spatial resolution Δ*x* = 0.001 *L* (green), staying constant within a wide range of field strength *E*. These resolution-related artifacts do not occur anymore for *R*_min_ (*E*), when the spatial resolution 0.0001 *L* (yellow) is chosen to be ten times higher.

**FIG. 13.**
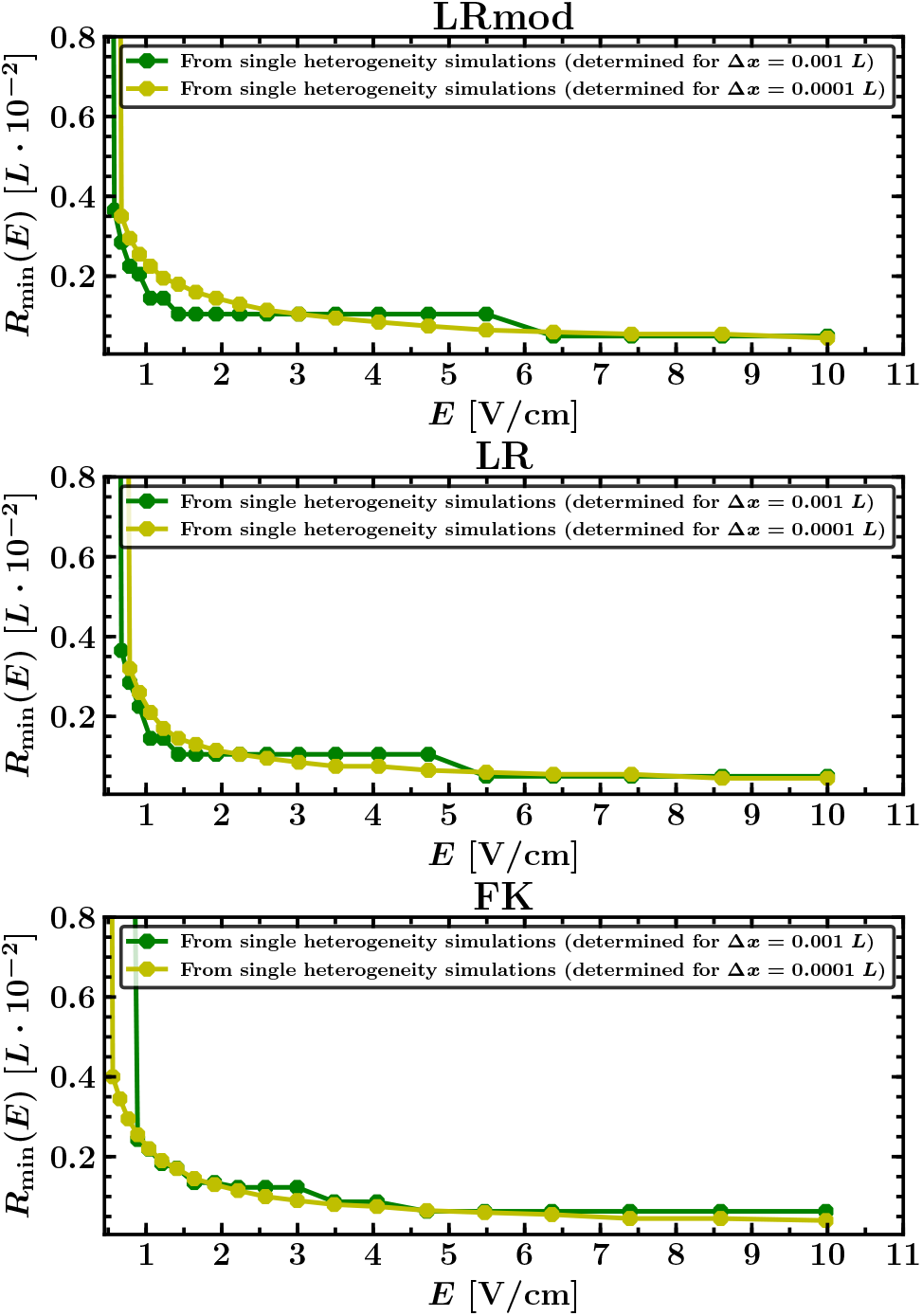
Minimal radius *R*_min_ (*E*) that is necessary for a heterogeneity to get activated by a pulse as a function of the electric field strength *E* of the pulse. The radii *R*_min_ (*E*) were determined in single heterogeneity simulations, where it was checked for each field strength *E* and a wide range of radii whether a circular heterogeneity that is placed in the center of the simulation domain gets activated or not by a pulse. For the spatial resolution Δ*x* = 0.001 *L* (green), which is used by most of the simulations in this thesis, resolution-related artifacts are clearly visible: *R*_min_ (*E*) decreases stepwise and not smoothly, staying constant within a wide range of field strength *E*, in contrast to *R*_min_ (*E*) determined for a ten times higher spatial resolution of 0.0001 *L* (yellow), where those resolution-related artifacts do not occur anymore.

Resolution-related artifacts may hence be expected to appear also for the density of activated heterogeneities *ρ* according to relation (A2). However, the relation (A2) does not take into account further resolution-independent factors like the current excitability of the surrounding tissue of a heterogeneity or whether there are other heterogeneities nearby that support or counteract the depolarization of the surrounding tissue. Such factors have also a decisive influence on whether a heterogeneity gets activated, and thus can compensate or at least mitigate the resolution-related factors considered in (A2). Indeed, for the activation density *ρ* determined directly as the average quotient of the number of activated heterogeneities after a pulse and the area size of excitable tissue right before the pulse, resolution-related artifacts are almost not observable, see figure 14. Moreover, figure 14 shows that the activation density *ρ* is independent from the fact whether it is determined in simulations where all heterogeneities with a diameter smaller than the grid spacing (*R <* Δ*x/*2 = 0.0005*L*) were set to the size of one pixel (blue), or whether it was determined in simulations where only heterogeneities with *R* ≥ Δ*x/*2 = 0.0005*L* were considered (cyan). In other words, the slight distortion of the size distribution of heterogeneties that stems from the relatively coarse discretization used in our simulation described above does not affect the activation density *ρ*.

**FIG. 14.**
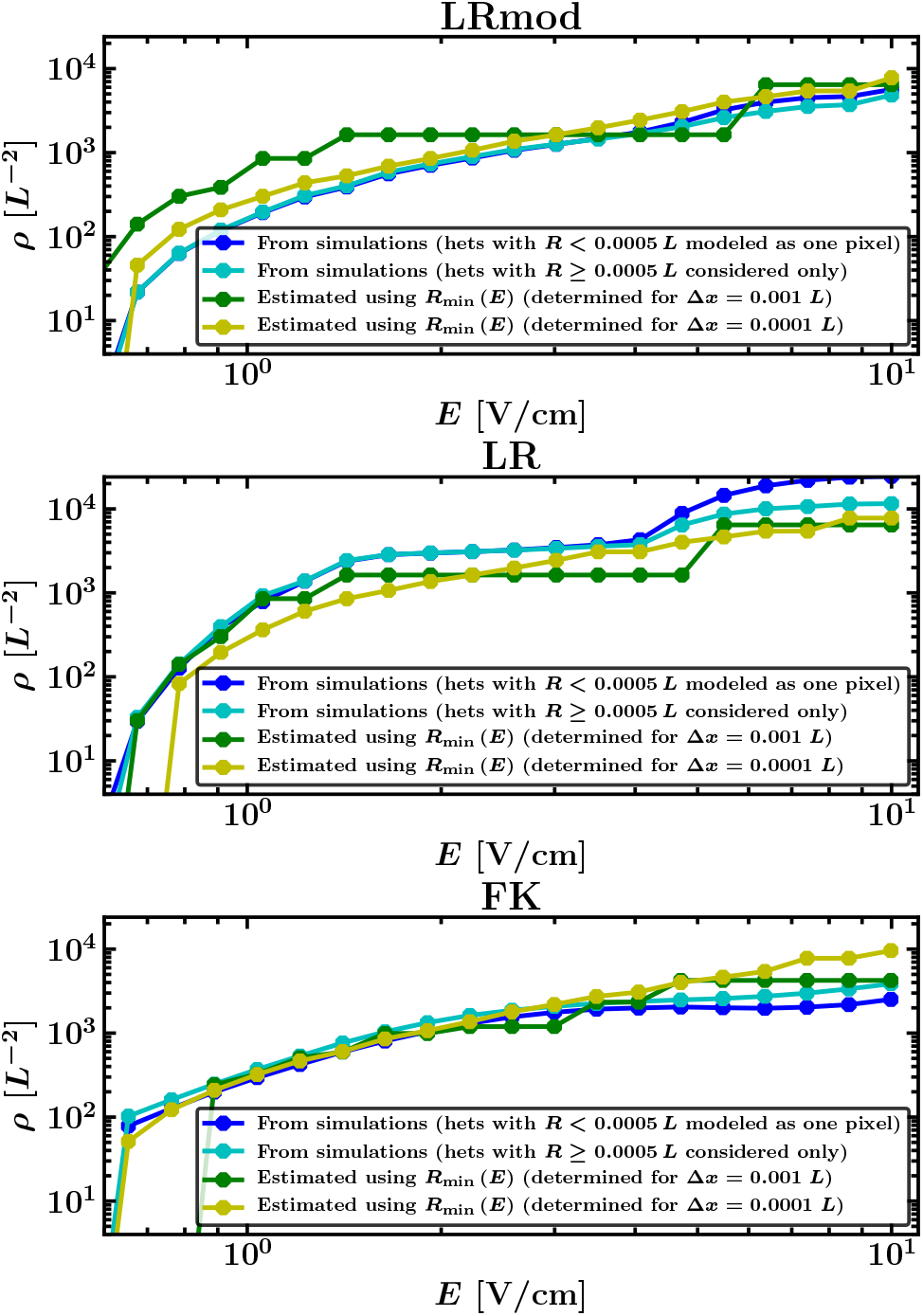
Density of activated heterogeneities *ρ*, as a function of the electric field strength *E. ρ* determined directly in simulations of each initial state of the three models (LRmod, LR, FK) as the average quotient of the number of activated heterogeneities after a pulse and the area size of excitable tissue right before the pulse, is plotted in blue for simulations where all heterogeneities with a diameter smaller than the grid spacing (*R <* Δ*x/*2 = 0.0005*L*) were set to the size of one pixel and in cyan for simulations where only heterogeneities with *R* ≥ Δ*x/*2 = 0.0005*L* were considered and placed on the simulation domain. Both curves are almost equal for field strength *E* below the single pulse defibrillation threshold (vertical dotted line) indicating that the chosen approach of setting heterogeneities with *R <* 0.0005*L* to the size of one pixel should not significantly impact the simulation results for LEAP. Furthermore, resolution-related artifacts due to the low spatial resolution of Δ*x* = 0.001 *L*, which occur for the minimal radius that is necessary for a heterogeneity to get activated *R*_min_ (*E*) (see Fig.13) and which could also be expected to occur for *ρ* (due to relation (A2)), are not observable: The course of both directly measured *ρ* (cyan and blue) is smooth and does not increase step-wise as one would expect at first, estimating *ρ* with equation (A2) via the minimal radius *R*_min_ (*E*) (green). Instead, the course is more similar to the estimation of *ρ* for a ten times higher spatial resolution of 0.0001 *L* (yellow), for which resolution-related artifacts of the underlying minimal radius *R*_min_ (*E*) do not occur anymore.

Furthermore it can be seen, that the approach how heterogeneities with a diameter smaller than the grid spacing are modeled has no significant impact on the density of activated heterogeneities *ρ*, at least not for field strength *E* below the single pulse defibrillation threshold, which were mainly used in this work. The course of *ρ* determined in simulations where all heterogeneities with a diameter smaller than the grid spacing were set to the size of one pixel (blue) is almost equal to the course of *ρ* determined in simulations where, in contrast, all these small heterogeneities were removed from the simulation domain (cyan).

Overall, it can be therefore assumed that the usage of the spatial resolution Δ*x* = 0.001 *L* should not significantly impact the simulation results in this work, since the density of activated heterogeneities *ρ*, which mainly characterizes the activation behavior of the heterogeneities, is neither significantly affected by the limitations of resolving heterogeneities correctly, nor by the modeling approach of setting heterogeneities with a diameter smaller than the grid spacing to the size of one pixel.

## Appendix B Macro-observables

### 1. Determination of the macro-observables

Fig.15 shows for an arbitrary chosen snapshot of the (LR) model (Fig.15a) exemplary the procedure for the determination of the refractory boundary length and other in this work and especially in Sec II C considered macro-observables. The procedure for the (LRmod) and (FK) model was the same.

**FIG. 15.**
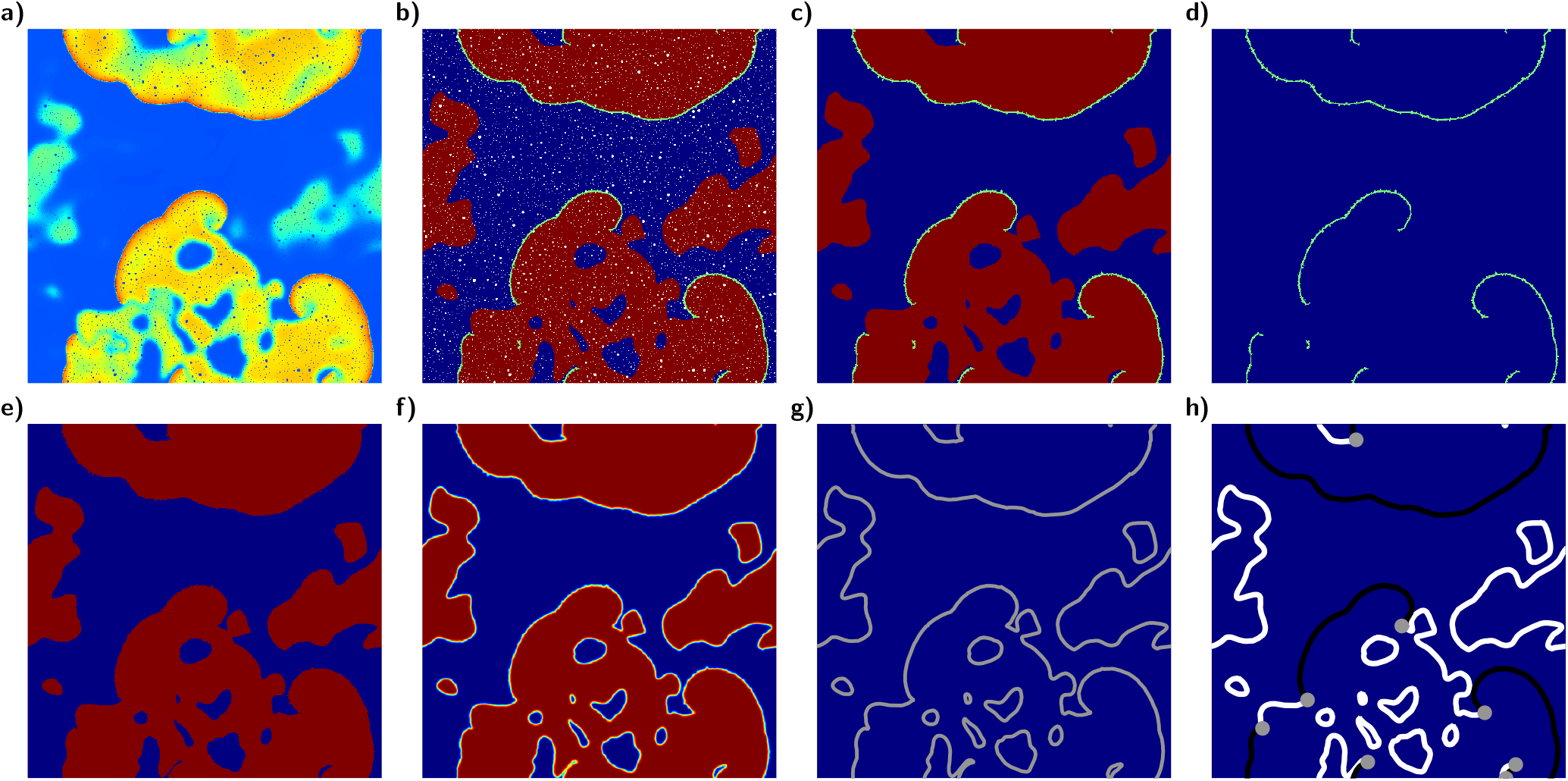
Segmentation of the refractory boundaries, excitation fronts and defects for an arbitrary chosen snapshot of the (LR) model. Figure (a) displays the transmembrane potential *V* in each point while (b) displays for all points whether they are excitable (blue), refractory (red), part of an excitation front (green) or part of a heterogeneity (white). As a first step, all heterogeneities were removed (c). Next the location of the front-points were saved (d) before all excitable points were set to a value of 0.0 (blue) and all other points to a value of 1.0 (red) (e). The application of a gaussian filter smoothed the edges (f) and enabled the determination of the borderlines between excitable and non-excitable parts of the tissue (grey) as the contour lines to a value of 0.5 (g) These borderlines were then segmented into excitation fronts (black) and refractory boundaries (white) (h) by comparing the location of these borderlines with the location of the front-points in (d). Defects (grey) were then simply determined as the connection points between the excitation fronts and refractory boundaries.

We have first assigned every point one of the following four categories (Fig.15b):

#### Heterogeneity (white)

These points are part of a heterogeneity and are simply given by the used distribution of conductivity heterogeneities.

#### Front (green)

These points are part of an excitation front and are characterized by a strong upstroke of the transmembrane potential after an excitation over the activation threshold and were determined by having a transmembrane potential greater than this activation threshold (*V > V*_thresh_) and a time derivative of the transmembrane greater than a certain appropriate limit value (*∂*_*t*_*V > V*_t,limit_) This thresholds and limits were given for the (LR-mod) and (LR) model by *V*_thresh_ = − 40 mV and *V*_t,limit_ = 10 mV*/*ms and for the (FK) model by *V*_thresh_ = −55 mV and *V*_t,limit_ = 5 mV*/*ms.

#### Excitable (blue)

These points are characterized by having a transmembrane potential lower than the activation threshold (*V < V*_thresh_) and releasing an action potential if the transmembrane potential exceeds this activation threshold. Therefore, an additional time integration had to be performed, where the transmembrane potential was set to the value of the activation threshold, and it was checked whether the transmembrane potential would achieve a value greater than 0.0 mV.

#### Refractory (red)

These points are simply given by the points which are not excitable and are further not part of a heterogeneity or an excitation front.

This representation of a state enabled us already to determine the fraction of excitable tissue as the number of excitable points divided by the total number of points without heterogeneities (*F*_Exc_ = *N*_Exc_*/*(*N*_total_ − *N*_Het_)). The number of excitable clusters *N*_cExc_ and the number of non-excitable(“refractory” or “front”) clusters *N*_cNExc_ were determined by a connected component labeling algorithm extended by the ability to consider periodic boundary conditions if necessary.

In order to determine the total length and number of all refractory boundaries (*L*_RB_ and *N*_RB_), the total length and number of all excitation fronts (*L*_Fr_ and *N*_Fr_) and the number of all defects (*N*_Def_), the segmentation of all excitation fronts, refractory boundaries and defects were needed. Therefore, first all heterogeneities had to been removed, which were done by simply turning iteratively all “heterogeneity”-points into “front” which are next to a “front”-point, the then remaining “heterogeneity”-points into “excitable” if they are next to an “excitable”-point, the then remaining “heterogeneity”-points into “refractory” if they are next to a “refractory”-point and again the then remaining “heterogeneity”-points into “front” if they are next to a “front”-point and so on until no “heterogeneity”-point were left (Fig. 15c). Next the location of the “front”-points were saved (Fig.15d) before all “excitable”-points were set to a value of 0.0 (blue) and all non-excitable-points(“refractory” or “front”) to a value of 1.0 (red), see Fig.15e. The application of a gaussian filter smoothed then the edges (Fig.15f) and enabled the determination of the borderlines between the excitable and non-excitable parts of the tissue (grey) as the contour lines to a value of 0.5 (Fig.15g). These borderlines were then segmented into excitation fronts (black) and refractory boundaries (white), see Fig.15g, by simply comparing the location of these borderlines with the location of the “front”-points (Fig.15d). Finally the defects (grey) were simply determined as the connection points between the excitation fronts and refractory boundaries.

The time derivatives *∂*_*t*_*F*_Exc_, *∂*_*t*_*L*_RB_ and *∂*_*t*_*L*_FR_ of the fraction of excitable tissue *F*_Exc_, the total length of all refractory boundaries *L*_RB_ and the total length of all excitation fronts *L*_FR_ were simply calculated as the difference quotients with the values of the previous time step.

#### 2. Analysis of the macro-observables

We tested all eleven in Sec II C introduced macro-observables, for all three models and in each case four different field strength *E*, see Table I. The values of the calculated Rank biserial correlation coefficients *r*_rb_ for all eleven macro-observables are listed in Table I. The strongest correlations with the defibrillation success *S* were found for the refractory boundary length *L*_RB_, which is additionally the only macro-observable at all that is highly correlated with *S* for all three models and field strength *E*. The refractory boundary length *L*_RB_ is therefore the most promising macro-observable to identify a common mechanism for defibrillation by periodic pacing in all three models.

Beside *L*_RB_, we found for the (LRmod) model six additional macro-observables that are comparable high correlated with *S*. These are given by the fraction of excitable tissue *F*_Exc_, the total length of all excitation fronts *L*_Fr_, the number of excitable clusters *N*_cExc_, the number of refractory boundaries *N*_RB_, the number of excitation fronts *N*_Fr_ and the number of defects *N*_Def_, see Table I. All these six have in common that they are highly correlated with the refractory boundary length *L*_RB_ within the (LRmod) model, see Table II. These correlations base on the fact, that the (LRmod) model exhibits stable spirals, which have roughly a typical characteristic size, width and curvature and where therefore typical size relations among these six macro-observables and *L*_RB_ exist, see Fig.3.

For the (LR) and (FK) model, these six macro-observables exhibit no strong correlation with successful defibrillation. These two models exhibit spatiotemporal chaos and unstable spirals, respectively, where these correlations do not or only in a weak manner exist. For these two models, we found with the time derivative of the fraction of excitable *∂*_*t*_*F*_Exc_ one additional macro-observable that is for at least three out of four considered field strength *E* comparable high correlated with *S*, see Table I. *∂*_*t*_*F*_Exc_ can be interpreted as the difference per time unit between the fraction of tissue that turns from non-excitable to excitable and the fraction of tissue that turns from excitable to non-excitable. These turnings from non-excitable to excitable take place at the refractory boundaries and vice versa from excitable to non-excitable at the excitation fronts. Thus, *∂*_*t*_*F*_Exc_ is proportional and therefore highly correlated to the difference between the total length of all refractory boundaries and excitation fronts (*L*_RB_-*L*_FR_), as shown in Table II. Therefore, it should feature only high correlations with the defibrillation success *S* if the correlations of *L*_RB_ and *L*_FR_ with *S* have opposite signs. That is the case for the (LR) and (FK) compared to the (LRmod) model and explains why this two models feature high correlations between *S* and the time derivative of the fraction of excitable tissue *∂*_*t*_*F*_Exc_ while the (LRmod) model does not.

Overall, macro-observables that have a strong correlation with the refractory boundary length *L*_RB_ itself, display also a good correlation with the defibrillation success probability. *L*_RB_ is altogether only tested macro-observable that allows a prediction of defibrillation outcome for all three models, which is a strong indicator that it might be a good candidate to decipher the underlying mechanism behind LEAP.

## Appendix C Snapshots of newly induced excitation fronts at the back of a plane wave for different field strength E

The in Sec III C 1 introduced density of newly induced excitation fronts *λ* (*E*) is usually field dependent. Fig.16 shows this field-dependency exemplary for the (LRmod) model at the back of a plane wave. The line density *λ* (*E*) or rather the number of excitation fronts that get induced at the back of this plane wave decreases with the electric field strength *E* and becomes zero for field strength *E* greater than the single pulse defibrillation threshold of 6 V*/*cm.

**FIG. 16.**
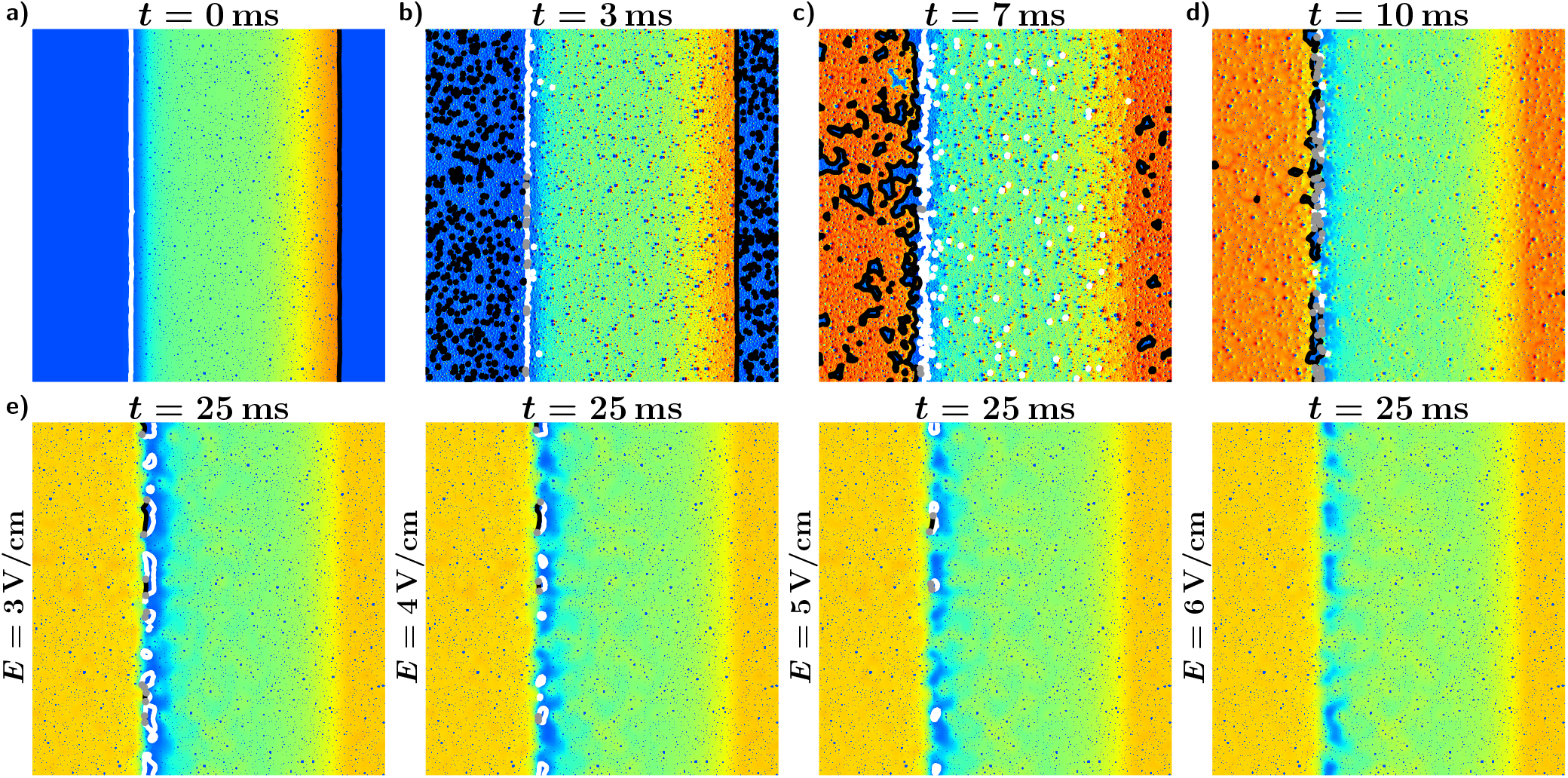
Displayed are for the (LRmod) model and different field strength *E* the snapshots of the transmembrane potential of the newly induced excitation fronts that arise right after a low-energy pulse at the back of a plane wave. Excitation fronts are plotted black, refractory boundaries are plotted white and the topological defects are marked as grey dots. (a) Plane wave right before the pulse. (b-d) Plane wave during the pulse when the pulse excites quickly the entire excitable tissue. (exemplary shown for a field strength of *E* = 3 V*/*cm) (e) Plane wave after the pulse, when almost the entire tissue is excited and only the newly induced excitation fronts are left. The number of newly induced excitation fronts *N*_Fr,induced_ decreases with the field strength *E* until *N*_Fr,induced_ will be zweo for field strengths *E* greater than the single pulse defibrillation threshold of 6 V*/*cm. Movies of the simulations are provided in the supplementary material.

## Appendix D Determination of the decay rate *k* and the line density of newly induced excitation fronts *λ*

In order to determine the in Sec III B introduced decay rate *k* (*E*) of the exponential relationship between termination probability *P* and refractory boundary length *L*_RB_ we first had to bin the data, since the termination probability *P* can only be determined as the fraction of successful termination events. Therefore, we sorted for each model and field strength *E* all the 10000 performed simulation runs by *L*_RB_, divided them into equally sized bins with a size of 200 and computed for each bin the termination probability *P* and the mean of the refractory boundary length *L*_RB_. The decay rate *k* (*E*) were then determined as the decay rate of the exponential regression between *P* and *L*_RB_, as visualized in Fig.5.

We determined the in Sec III C 1 introduced line density of newly induced excitation fronts *λ* (*E*) as the average ratio between the number of newly induced excitation fronts right after the pulse and the refractory boundary length *L*_RB_ right before the pulse of the 10000 simulation runs that were performed for each model and electric field strength *E*. The right choice of the time *t*_Nind_ after the pulse for counting the newly induced excitation fronts is a balance act between counting to early, before the entire excitable tissue got excited and when therefore more than the looked for newly induced excitation fronts are left, and counting to late when some of the newly induced excitation fronts have already fused. Especially for low field strengths *E*, the time window for counting the right number of newly induced excitation fronts is small if not non-existent, since the pulse needs more time to excite the entire excitable tissue and moreover the density of newly induced excitation fronts *λ* (*E*) along the refractory boundaries is high and those fronts start earlier to fuse. In order to enlarge this time window towards lower times, only excitation fronts were counted that move or that are directly or indirectly connected with fronts or refractory boundaries that move into the progressively parts of the tissue. All other fronts die out in this case anyway when the entire excitable tissue got excited. Therefore, it was checked whether an excitation front sits on a contour line (check out Appendix B 1 for more details about the segmentation of the refractory boundaries and excitation fronts) that partially lays on tissue that was right before the pulse not excitable. However, this procedure does not work completely fault-free for very low times *t*_Nind_ and thus this time window remains still small.

By experience from looking at a lot of LEAP episodes, we found, that counting of the newly induced fronts works best at a time *t*_Nind_ = 25 ms for the (LRmod) and (FK) model and *t*_Nind_ = 50 ms for the (LR) model. These times *t*_Nind_ lay within this time window, where the right number of newly induced fronts get counted, as we will show in the following, and have been therefore used in this work for the determination of the line density of newly induced excitation fronts *λ*.

In Fig.17 is plotted *λ* in dependence of the field strength *E* for different times *t*_Nind_ at which the corresponding numbers of newly induced excitation fronts were counted. The looked for time window can be identified in this plot by the range of times *t*_Nind_ where those courses of *λ* (*E*) are equal or at least very similiar, since the right number of newly induced excitation fronts and thus the right *λ* does not depend on *t*_Nind_. For the (LR-mod) and the (FK) model, that is the case for *t*_Nind_ between 20 ms and 25 ms. For higher *t*_Nind_, the courses become more flatten towards low field strength *E*, since the fusing of the newly induced excitation fronts increasingly becomes an issue, while for lower *t*_Nind_ the courses become steeper due to additional fronts that exist at this early times after a pulse and were wrongly counted. For the (LR) model such a time window does not exist. Nevertheless, the curves of *λ* (*E*) are very close together for *t*_Nind_ between 40 ms and 70 ms and the most accurate time for counting the number of newly induced excitation fronts for the (LR) model might lay in this range.

**FIG. 17.**
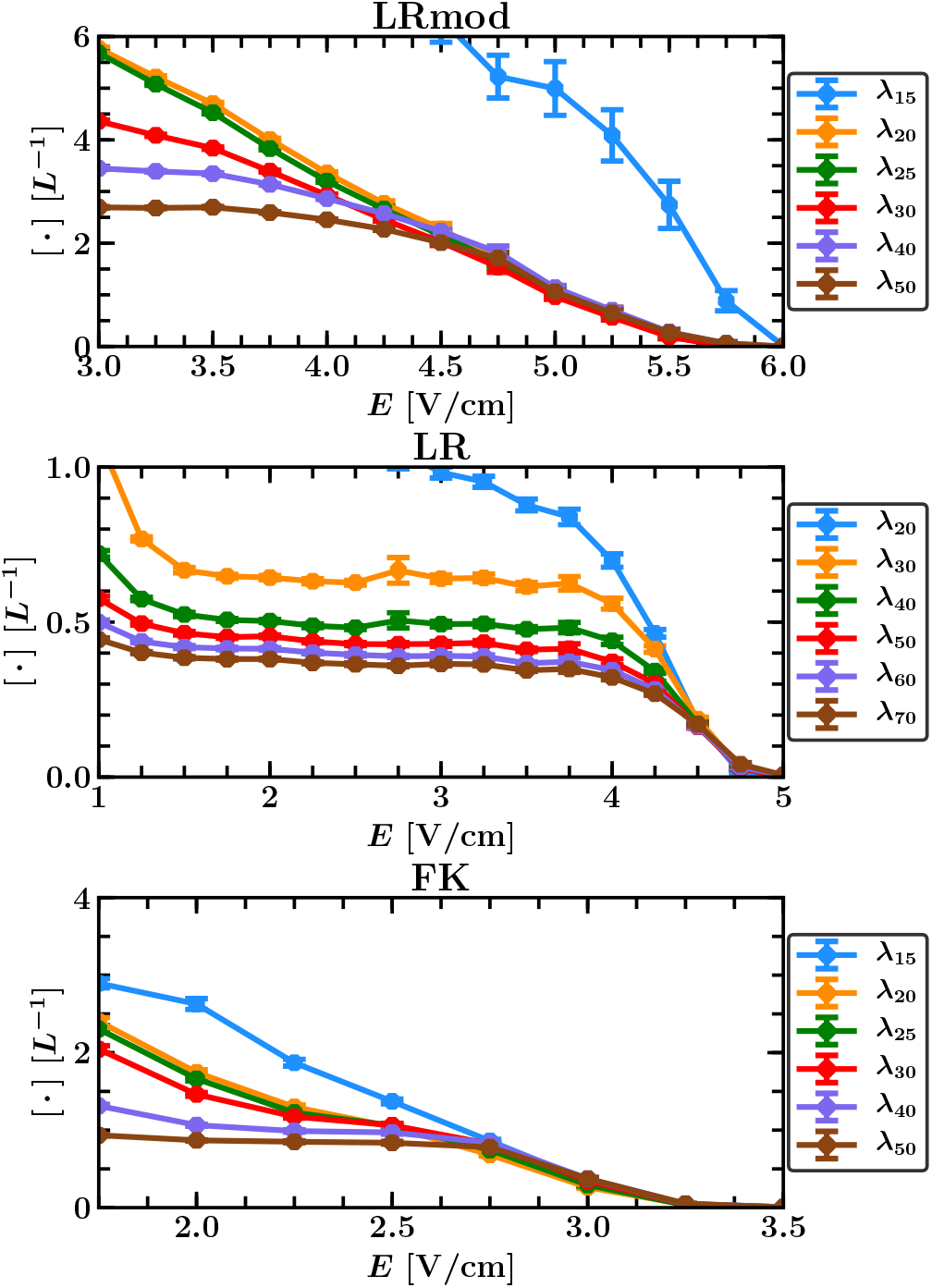
Line density of newly induced excitation fronts *λ*_*T*_ as a function of the electric field strength *E* determined as the average fraction between the number of newly induced excitation fronts, counted at a time *T* after the pulse, and the refractory boundary length *L*_RB_ right before the pulse.

## Notes

### Competing Interest Statement

The authors have declared no competing interest.

